# Methylation dynamics in the decades preceding acute myeloid leukaemia

**DOI:** 10.1101/2025.06.26.661643

**Authors:** Adriana V.A. Fonseca, Christopher Boniface, Caroline J. Watson, Samuel Hackett, Calum Gabbutt, Akemi D. Ramos-Yamasaki, Amanda B. Tan, Jose Montoya, Ruslan Strogantsev, Lars L. P. Hanssen, Yuexuan Zhang, Sophia Apostolidou, Aleksandra Gentry-Maharaj, Sadik Esener, Paul Spellman, Trevor Graham, Usha Menon, Hisham Mohammed, Jamie R. Blundell

## Abstract

DNA methylation is emerging as a highly sensitive and specific marker of cancer initiation and progression. How these cancer-specific methylation changes are established in the decades before cancer, however, remains largely unknown. Here, we use a unique collection of longitudinal blood samples collected annually up to 15 years prior to a diagnosis of acute myeloid leukaemia to sensitively track the dynamics of DNA methylation changes at high temporal resolution. We identify thousands of differentially methylated regions (DMRs) that exhibit altered patterns of methylation up to 10 years before acute myeloid leukaemia (AML) diagnosis. Most of these DMRs are strongly associated with expanding clones carrying somatic driver mutations. We identify a subset of ‘epigenetic driver’ DMRs characterised by recurrent CpG alterations that are highly shared across pre-AML cases. These are likely to reflect early epigenetic reprogramming associated with AML development. We also reveal large numbers of stochastic ‘passenger’ CpGs whose differential methylation results from hitch-hiking with clonal expansions driven by somatically acquired genetic events. These passenger CpGs can be exploited for lineage tracing to discover clonal expansions driven by missing driver mutations. Our findings show widespread changes in methylation patterns during the early stages of cancer development which could be utilised for risk prediction and therapeutic intervention.

## Introduction

Cancer emerges through an evolutionary process that involves the acquisition of genetic and epigenetic changes that co-opt programs of tissue homeostasis, differentiation, and repair ^*1–3*^. Although genetic drivers of cancer are known to be acquired many years or even decades before cancer diagnosis ^*4–9*^, much less is known about when epigenetic alterations arise and how they influence cellular fitness ^*10*^. DNA methylation is a dynamic epigenetic mark that plays a crucial role in controlling cellular fate and identity during development ^*11*^, differentiation ^*12*^, ageing ^*13*^ and carcinogenesis ^*14*^. Cancer-specific changes in DNA methylation are observed in a wide range of cancers ^*15,16*^, and fall into two broad categories: hypo- and hypermethylation. Genome-wide hypomethylation can lead to loss of transcriptional repression, particularly at repetitive elements, contributing to genomic instability and promoting cellular transformation. ^*15,17,18*^. Hypermethylation typically occurs at specific CpG islands, particularly within the promoters of tumour suppressor genes, leading to reduced gene expression ^*19–21*^. This epigenetic alteration can contribute to tumourigenesis by disrupting cancer-related pathways ^*20,22–25*^.

Beyond their role in advancing our understanding of cancer biology, methylation changes offer a promising avenue for improving cancer risk prediction and early detection. Cell-free DNA methylation patterns have been used to diagnose the presence and type of stage I/II cancers with high sensitivity ^*26*^, and breast and ovarian cancer-specific signatures are detectable in samples collected years before diagnosis ^*27–30*^. However, the ‘life history’ of cancer-specific methylation changes remains largely unknown. When are patterns of altered methylation first established? What is the relationship between methylation change and expansion of genetic driver alterations? Which methylation changes constitute ‘epigenetic drivers’ implicated in disease biology, and which are ‘epigenetic passengers’ hitch-hiking on clonal expansions? Answers to these questions could improve our understanding of the programs co-opted in the earliest stages of cancer, open up avenues for improving early cancer detection, and point to therapeutic targets.

The blood cancer acute myeloid leukaemia (AML) represents a model cancer for addressing these questions. Many of the early genetic drivers of AML occur in epigenetic regulators such as *DNMT3A, TET2*, and *IDH1/2*^*31,32*^. Methylation profiles of leukaemic cells are distinct from those of healthy haematopoietic stem cells (HSCs), from which the disease originates ^*33–36*^. Furthermore, methylation patterns have been used to define disease subtypes ^*33,37–42*^ and are predictive of patient prognosis ^*33–35,38,40,43,44*^. Hypomethylating agents such as azacitidine and decitabine are commonly used as AML treatments, particularly in older patients ^*45–47*^. Whilst methylation differences between samples taken over a decade before diagnosis of AML and controls have been identified ^*48*^, little is known about their dynamics. We hypothesised that analysing serial samples collected over a ten-year period from individuals who later developed AML, alongside age-matched controls, could uncover early epigenetic changes specifically linked to future cancer development. By tracking methylation changes over time and integrating these patterns with longitudinal profiling of somatic variants from the same samples, we aimed to elucidate the relationship between somatic driver mutations and epigenetic alterations in the years preceding a cancer diagnosis.

## Results

### Whole-genome discovery of differential methylation

We first sought to identify regions that exhibit differential methylation (differentially methylated regions, DMRs) during AML development at whole-genome scale (Fig. 1). We analysed whole genome bisulphite sequencing (WGBS) data from the BLUEPRINT consortium ^*50*^ comprising purified AML blasts (n=21) and healthy HSCs (n=8). To sensitively detect regions with robust methylation differences despite limited sequencing depth, we developed a statistical method tailored specifically for low-depth WGBS data which identifies regions of different sizes with large statistical differences between AML and HSC samples (Fig. 1a, Methods).

**Fig. 1.**
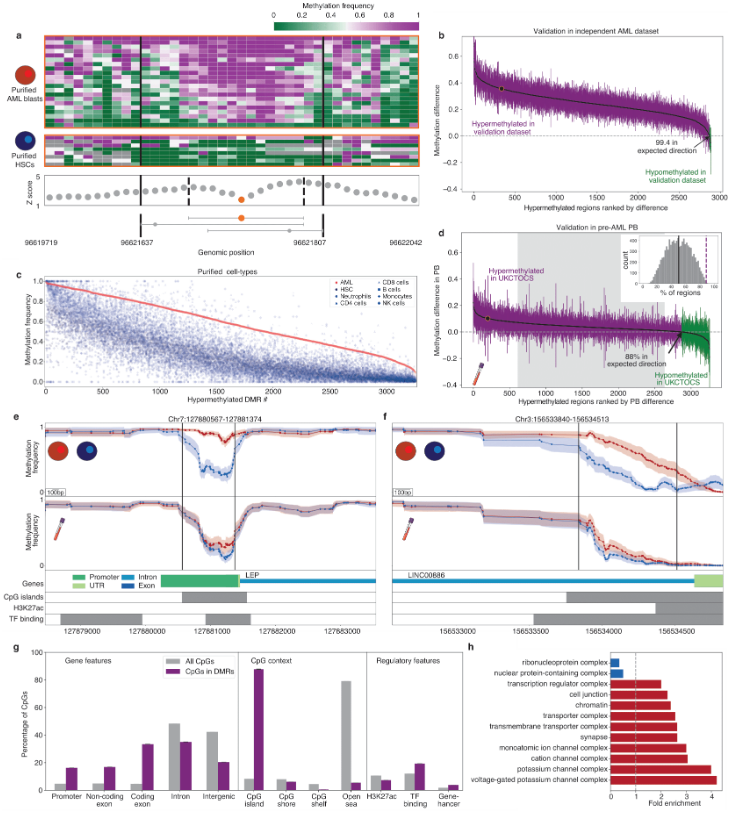
DMR discovery and validation. a. Example of DMR discovery. Seed CpGs (orange point) grow to incorporate neighbouring CpGs creating contiguous regions where statistical differences in methylation frequency between AML blasts (red) and healthy HSCs (blue) is maximized (dashed lines). Overlapping regions are merged into one DMR (black lines). **b**. Mean methylation difference (black points) between an independent cohort of AML samples from Glass *et al*. ^*49*^ and BLUEPRINT ^*50*^ HSC samples, for all hypermethylated regions identified from our discovery. Coloured bars show standard deviation and direction of difference. **c**. Mean methylation level of each DMR in AML blasts (red points), HSCs, and multiple peripheral blood cell types (blue points). **d**. Mean methylation difference between peripheral blood (PB) samples from pre-AML cases and matched controls (black points), for all hypermethylated regions. Coloured bars show standard deviation and direction of difference. Inset: Distribution of % of regions in the expected direction from permuted data (grey) compared to the observed % of regions in the expected direction (purple dashed line, p<0.01). Grey shading reflects 95% confidence intervals from the permutation analysis. Highlighted region (orange) in panels **b** and **d** corresponds to the region in panel **a. e, f** Example DMRs in regions with (**e**) or without (**f**) previous support for AML-association. Moving average methylation levels in AML-blasts (top panel, red) versus HSCs (top panel, blue). Moving average methylation levels in the same region from peripheral blood (middle panel) collected from pre-AML cases (red) and healthy controls (blue). Shading represents standard error on the mean. (Bottom panels) Gene isoforms, CpG islands, ChIP-seq reads relating to H3K27 acetylation in K562 cells, transcription factor binding in K562 cells plotted for the region in question. **g** Fraction of all CpGs (grey) or CpGs within validating DMRs (purple) for the labelled feature. **h** Fold enrichment for GO terms found to be significantly enriched or purified for the set of genes overlapping the validating DMRs.

This method identified approximately 70,000 hypomethylated and 3,200 hypermethylated DMRs in AML compared to HSCs (Extended Data Fig. 1, Methods). Over 99% of these DMRs showed consistent methylation directionality in an independent dataset comprising 119 AML samples ^*49*^ (Fig. 1b and Extended Data Fig. 2, methods). To further validate the AML-specificity of these DMRs, we compared their methylation profiles in AML samples to those from various healthy peripheral blood cell types. Hypermethylated DMRs were highly AML-specific (Fig. 1c), whereas hypomethylated regions tended to reflect HSC-specific differentiation signatures (Extended Data Fig. 3), aligning with prior observations of differentiation-associated methylation ^*12,51*^.

We hypothesised that these hypermethylated DMRs might reflect early events in AML pathogenesis which could be detected in peripheral blood. To test this, we performed WGBS on peripheral blood samples collected within the two years prior to AML diagnosis from the UK Collaborative Trial of Ovarian Cancer Screening (UKCTOCS ^*52,53*^) cohort. This study stored serum which contained large quantities of genomic DNA derived from granulocytes (Extended Data Fig 4, Methods). We confirmed significant enrichment of hyper-methylation at these DMRs in pre-AML blood samples compared to matched controls (88% validation, permutation test p < 0.01, Fig. 1d). As expected, this was not the case for hypomethylated regions (Extended Data Fig. 5). These findings strongly support the potential for hypermethylated DMRs to be early biomarkers of AML development.

Our analysis uncovered regions previously associated with AML progression and prognosis, e.g. LEP ^*54,55*^ (Fig. 1e, Supplementary Table 1) as well as novel loci, e.g. a number of long non-coding RNAs (Fig. 1f). The majority of AML-specific DMRs occurred in CpG islands and showed strong enrichment for promoters, exons and transcription factor binding motifs (Fig. 1g). Gene ontology analysis of the *∼* 1800 genes that overlap with the DMRs defined showed strong enrichment for transcriptional regulation, chromatin remodelling and neurodevelopmental genes (Fig. 1h). Taken together, our findings highlight thousands of AML-specific DMRs whose methylation dynamics could be exploited for early detection and risk prediction.

### Longitudinal DNAme sequencing of pre-AML bloods

To quantitatively measure the dynamics of DNA methylation changes during the earliest stages of cancer initiation, we performed deep targeted methylation sequencing of 265 longitudinal blood samples collected annually from 50 women who later developed AML and 50 age-matched controls, spanning up to 11 years (Fig. 2a, and Extended Data Fig. 6). We designed a custom hybrid capture panel (Twist Biosciences) targeting *∼*5,000 regions, comprising DMRs identified in our whole-genome discovery analysis described above, regions showing differential methylation in peripheral blood, CpGs associated with epigenetic ageing ^*56*^, and regions of high methylation entropy ^*57*^ (Methods). In total our panel targets *∼*150,000 CpGs across 2.1Mb (Fig. 2b, Extended Data Fig. 7, Methods). Using enzymatic methylation sequencing ^*58*^ with high DNA input (30ng), we achieved high unique molecular coverage (*∼*750x per CpG site per sample), providing unprecedented temporal and frequency resolution for detecting subtle longitudinal changes in methylation levels (Fig. 2c-f).

**Fig. 2.**
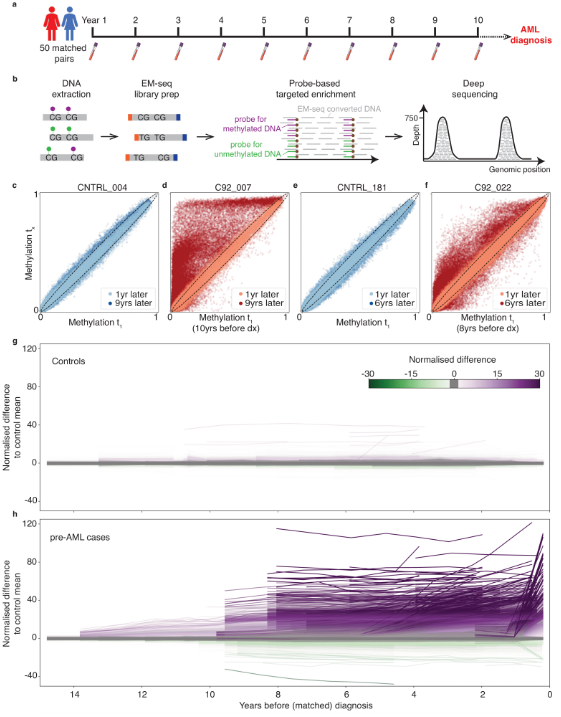
Deep targeted sequencing of longitudinal samples from healthy and pre-malignant blood. a. Longitudinal blood samples collected annually from 50 future AML cases and 50 age-matched controls from the UKCTOCS cohort. **b**. Workflow for deep methylation sequencing targeting *∼*5,000 AML-specific DMRs. **c-f**. Scatter plots of methylation levels of all *∼*150,000 CpGs at first (x-axis) and subsequent (y-axis) time points in two cases and matched controls. Dashed line shows expected (2*σ*) variation from technical noise due to finite molecular coverage (here, 300x). **g, h** Methylation frequency trajectories, relative to controls, of each of the *∼*5,000 DMRs in each of the 50 controls (**g**) and pre-AML cases (**h**) standardized by the standard deviation observed in all controls (Methods).

This high-resolution longitudinal dataset reveals striking differences in epigenetic evolution in pre-AML cases compared to controls. DNA methylation is remarkably stable in controls across multiple years and highly consistent across individuals (Fig. 2c,e,g). In stark contrast, pre-AML cases exhibit widespread, progressive methylation changes, detectable more than a decade before clinical diagnosis (Fig. 2d,f,h). The consistent increases in hypermethylation at thousands of loci indicate that substantial epigenetic reprogramming is an early feature of AML evolution rather than a late consequence of cancer progression (Fig. 2h and Extended Data Fig. 8). These early epigenetic alterations occur extensively across diverse genomic contexts, suggesting a widespread reconfiguration of the epigenetic landscape preceding overt malignancy, highlighting their potential as early biomarkers.

### Association between methylation and genetic drivers

To explore the relationship between methylation dynamics and underlying genetic events, we combined our deep methylation data with sequencing of somatic mutations in AML-associated genes from the same longitudinal samples using a highly sensitive duplex error-corrected DNA sequencing approach previously reported ^*59*^. We then assessed whether specific methylation changes could be statistically linked to particular somatic clones by identifying CpGs whose methylation level through time was strongly correlated with the dynamics of the somatic clone (Fig. 3a-d, Methods).

**Fig. 3.**
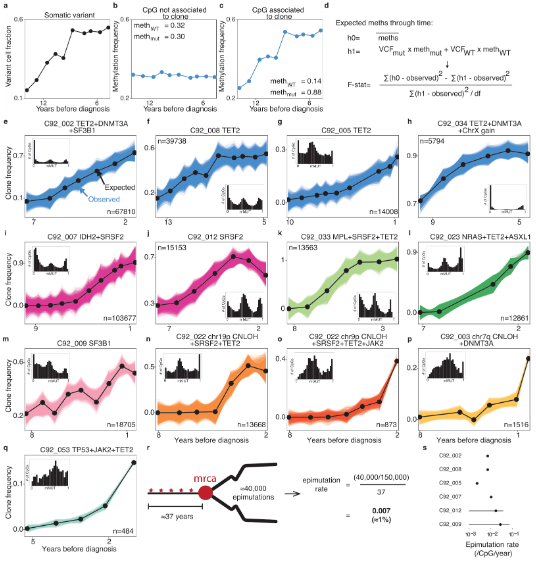
Association between CpG methylation and somatic clone trajectories. **a** Example trajectory for a somatic variant (TET2 p.G1361V in C92_008). **b-c** Methylation frequency through time of example individual CpG sites in the same donor. **d** Alternative hypotheses for expected methylation through time, and f-statistic calculation to test the significance of model improvement under h1 compared to h0. **e-q** Observed variant frequency trajectories of somatic driver variants (black data points) and the clone frequencies implied by individual CpGs (coloured lines) whose methylation frequency trajectories strongly associate with the variant frequency trajectory. The variant associated with is in the first gene mentioned in the title: where the variant is within a larger clone, other mutated genes in that clone are stated. Inset: distribution of inferred variant clone methylation levels for all CpGs associated to the clone. The number of CpGs significantly associated to the variant is also shown. For e-o FDR<0.05, for p-q FDR<0.1. **r** Schematic of epimutation rate calculation in example donor (C92_008) **s** Estimated epimutation rate in each donor, based on number of changes from WT background (number of associated CpGs) and inferred clone establishment times ^*59*^.

We found strong associations between specific somatic clones and tens of thousands of CpGs exhibiting altered methylation trajectories (range: 484–103,677 CpGs per clone, Fig. 3e-q), including for mutations not classically associated with epigenetic regulation. Notably, methylation levels associated with these expanding clones typically formed discrete peaks around methylation fractions of 0, 0.5, and 1 (insets), consistent with inheritance from the single ancestral cell initiating each expansion ^*60,61*^. This pattern suggests that the majority of methylation alterations represent stable epigenetic marks inherited from the founder cell of each clonal expansion.

We next leveraged these data to quantify the rate of spontaneous methylation changes (‘epimutation’) occurring within HSC lineages (Fig. 3r, Methods). By integrating the number of altered CpGs with estimates of the age at which clones arose ^*59*^, we inferred an average epimutation rate in pre-leukaemic HSCs of between 0.1%–1% per CpG per year across our panel (Fig. 3s), in line with previous estimates ^*61*^. This epimutation rate, approximately six orders of magnitude higher than somatic mutation rates ^*62*^, creates extensive epigenetic diversity across HSC lineages.

### Recurrent epigenetic drivers in pre-AML

Next, we sought to systematically identify DMRs recurrently altered in pre-AML cases but never altered in controls, as these could represent early, functional ‘epigenetic drivers’ implicated in AML initiation (Fig. 4). We began by identifying all CpGs in a sample that were statistical outliers relative to control methylation profiles (Fig. 4a and Extended Data Fig. 9, methods). Pre-AML cases generally exhibited higher epimutation burdens than controls, and epimutation burden was significantly associated with the presence of a somatic clone at >5% VAF (Mann-Whitney U test, *p* = 2.16 *×* 10^*−*16^, Fig. 4b). However, some donors displayed low epimutation burdens despite detectable somatic mutations (e.g. C92_019), whereas others showed high epimutation burden without detectable somatic mutations (e.g. C92_052), highlighting that we could be missing important genetic and epigenetic events in our targeted panels.

**Fig. 4.**
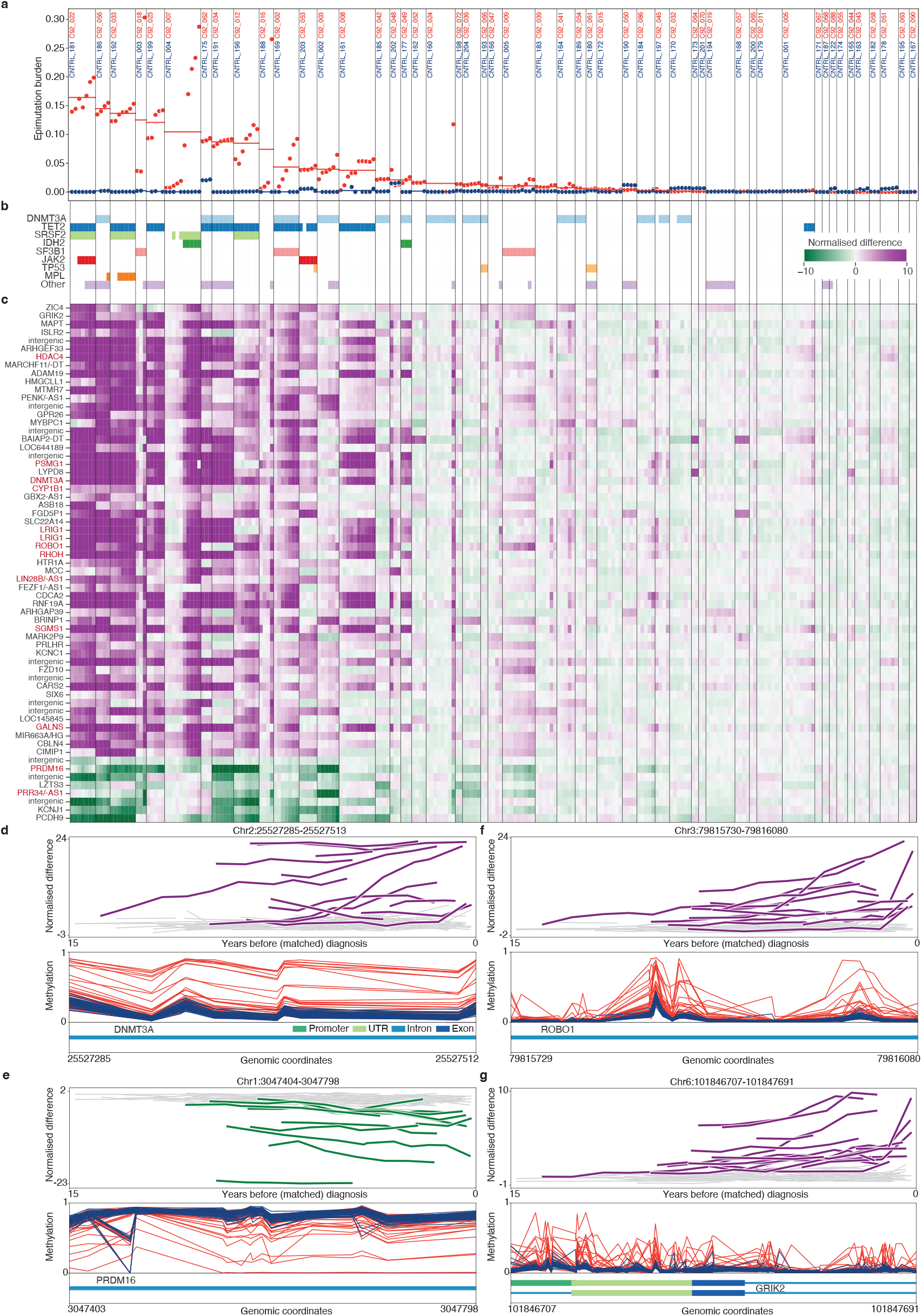
Methylation dynamics in healthy and pre-malignant blood. **a** Proportion of CpGs in each sample for which the absolute *z*-score is above 6. Each matched pair is separated by black lines, with mean epimutation burden shown for the case samples (red) and control samples (blue). **b** Presence of somatic change above 5% VAF, for each case sample. **c** Heatmap showing region *z*-score (mean of all CpGs in region) across regions (rows) and case samples (columns). Regions shown were never changed in controls, and either hypermethylated in 14 or more cases or hypomethylated in 10 or more cases. Genes highlighted by red text have been previously linked to AML biology. Where labelled as <gene>/-AS1, both the gene and its antisense transcript intersect the region. **d-g** Example regions. (Top panel) Mean region *z*-score through time in all donors. Cases changed highlighted by coloured lines. (Middle panel) Mean methylation for each CpG in the region, for the last timepoint in each case donor (red) and control donor (blue). (Bottom panel) Genomic context.

To identify candidate driver regions, we defined DMRs as AML-specific if more than 10% of CpGs within the region were significantly altered in a consistent direction across multiple (*>*5) pre-AML donors but not in any controls (Fig. 4c and Extended Data Fig. 10, Methods). This approach identified approximately 1,600 AML-specific regions that were recurrently altered in up to 40% of pre-AML donors (permutation test *p <* 0.05, Extended Data Fig. 11).

Many of these putative epigenetic driver regions overlapped genes with well-established roles in AML pathogenesis. These included the frequently mutated epigenetic regulator *DNMT3A* ^*32*^ (Fig. 4d), as well as *PRDM16* (Fig. 4e), a transcription factor critical for HSC maintenance whose increased expression and altered methylation have been previously associated with AML progression and prognosis ^*63–67*^. Another notable example is *ROBO1* (Fig. 4f), a receptor involved in axon guidance and neural migration, for which there is a growing body of evidence of a tumour-suppressor role in AML ^*68–70*^.

Strikingly, our analysis revealed that ROBO1 was just one of many recurrently altered epigenetic driver regions linked to neurodevelopmental processes (Extended Data Fig. 12, Supplementary Table 1). This unexpected enrichment included genes involved in axon guidance and neuronal connectivity (e.g., ROBO1, SLIT2, SEMA4C), synaptic signalling (e.g., GRIK2, PENK), and neuronal differentiation and central nervous system patterning (ZIC4, ISLR2). For instance, highly recurrent hypermethylation was observed in GRIK2 (18 cases, Fig. 4g), encoding a glutamate receptor involved in synaptic signalling, previously implicated through hypermethylation in gastric ^*71,72*^, lung ^*73*^, bladder ^*74,75*^ and oral ^*76,77*^ cancers. Together, these findings suggest that aberrant methylation of neurodevelopmental pathways could play an important yet under-explored role in early leukaemogenesis.

### CpG lineage tracing for clonal expansion discovery

Finally, we explored whether certain CpGs could serve as heritable epigenetic markers (‘passenger’ epimutations) to identify clonal expansions driven by genetic alterations outside our targeted panel of canonical AML genes. Such clonal expansions of unknown genetic origin have previously been described in clonal haematopoiesis ^*78,79*^. To identify suitable CpG markers, we focused on regions that are unlikely to be subject to active epigenetic regulation, yet stable enough to serve as markers of cell lineage.

We selected four samples where large clonal expansions had been identified by somatic sequencing (‘positive’) and four samples without a detectable clonal expansion (‘negative’). Adapting previous approaches ^*60,61*^, we identified a class of CpGs termed ‘fluctuating CpGs’ (fCpGs) that fulfilled three criteria: (i) were consistently polymorphic (methylation frequency near 0.5) in negative samples, (ii) became clonal in positive samples, and (iii) exhibited methylation states clonally fixed in either direction (fully methylated or unmethylated) across different positive samples (Methods). This approach yielded approximately 1,000 fCpGs sufficiently covered by our sequencing panel (Supplementary Table 2).

In polyclonal populations (i.e. those without significant clonal expansions), the frequency distribution of fCpGs is expected to be unimodal, centred around 0.5, reflecting diverse cellular ancestry dating back to embryonic development ^*80*^ (Fig. 5a). Conversely, in samples experiencing recent clonal expansions, the frequency distribution develops a characteristic trimodal or ‘W-shaped’ pattern, capturing the methylation state of the single ancestral cell initiating the expansion ^*60*^.

**Fig. 5.**
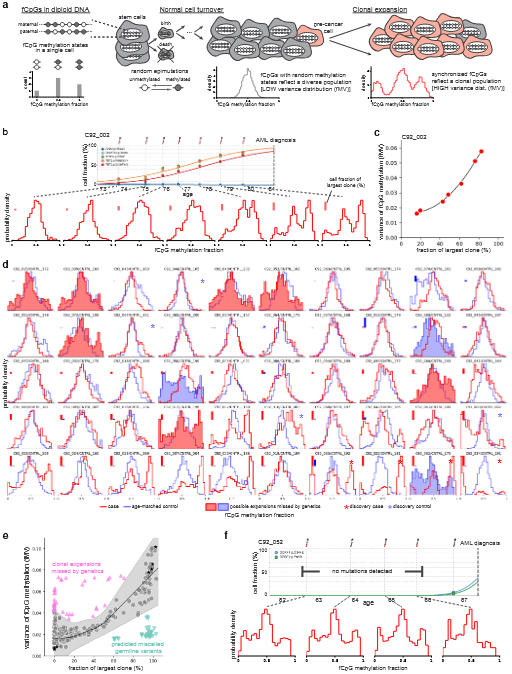
CpG lineage tracing for clonal expansion discovery. a. Using fCpGs for detecting clonal expansions. Populations without a recent common ancestor (grey) become uncorrelated producing a single peak in the fCpG distribution. Clonal expansions (red) introduce strong correlations giving rise to a trimodal feature in the distribution. **b**. The emergence of a ‘W’ feature in the fCpG distribution associated with a clonal expansion detected from somatic alterations. **c**. The measured variance in the fCpG distribution (fMV) as a function of the largest clonal cell fraction (red points) follows the expected quadratic relationship (*R*^2^=0.997, line). **d**. fCpG distributions for all 50 final time point samples in pre-AML cases (red) and controls (blue). Largest somatic clone detected is denoted by a solid bar on each panel. Samples used in fCpG discovery denoted by an asterisk. Filled distributions highlight expansions captured by fCpGs that were missed by genetic sequencing. **e**. fCpG distribution variance as a function of cell fraction for all samples. 11-fold expansion of 95% confidence intervals (grey shading) generated from a polynomial regression (black line) fit to data from three donors with expanding clones were used to identify outlier libraries (Extended Data Fig. 14, Methods). Pink (△) indicates samples where the fMV suggests the presence of an expansion missed by genetics. Teal (▽) indicates likely germline variants. Black stars indicate fCpG discovery libraries (asterisked in d). **f**. Example of an early somatic sweep missed by genetic sequencing.

To validate the utility of fCpGs for detecting clonal expansions, we analysed methylation distributions through time in samples with known expanding clones, using somatic mutations as the ground truth (Fig. 5b, Extended Data Fig. 13). We demonstrated that the expansion of a quadruple mutant clone (*DNMT3A*, biallelic *TET2, SF3B1*) clearly generates the characteristic ‘W-shaped’ methylation distribution as the clone expands. Furthermore, we observed a quantitative relationship between clonal cell fraction and variance in the fCpG distribution (Fig. 5c). Our analysis indicates that fCpG-based lineage tracing is capable of reliably identifying clonal expansions with cell fractions above approximately 20%.

We next analysed the fCpG methylation frequency distributions across all final time points from pre-AML cases and matched controls (Fig. 5d). Among controls, 44 of 47 donors where no large somatic clones were identified by targeted sequencing (94%) exhibited no detectable ‘W’ feature. However, in 3 control donors, we observed increased variance in the fCpG distribution beyond what would be expected based on detected clonal expansions alone (Fig. 5d histograms shaded blue, Fig. 5e, Extended Data Fig. 14, Methods). These findings suggest that broader genetic sequencing in these donors would likely uncover previously undetected clonal haematopoiesis driven by a clone with cell fraction >20%. Similarly, among pre-AML cases, of the 26 donors with a detectable somatic mutation above 20% cell fraction, 22 (85%) displayed the expected ‘W’ distribution. The remaining 4 cases likely represent miscalled germline variants. We also identified five pre-AML donors without detectable somatic mutations who nevertheless exhibited clear ‘W’ features (Fig. 5d histograms shaded red, Fig. 5e), suggesting these may carry cryptic clonal expansions driven by mutations outside our sequencing panel. Some of these expansions were already near fixation at the earliest sampled time points, years before clinical AML diagnosis (e.g. C92_052, Fig. 5f, Extended Data Fig. 15). Together, our findings confirm the utility of fCpG-based lineage tracing as an unbiased and potentially cost-effective method for detecting and characterising early clonal expansions.

## Discussion

In this study, we identified thousands of genomic regions exhibiting differential methylation during AML development. Hypermethylated DMRs, identified through comparison of AML blasts and healthy HSCs, were highly specific for AML, in contrast with hypomethylated regions, which predominantly reflect differentiation patterns. Remarkably, these AML-specific methylation signatures were stable enough to be reliably detected in DNA derived from peripheral blood, years before AML diagnosis.

Leveraging deep targeted methylation sequencing of longitudinal blood samples collected annually over approximately a decade leading up to AML diagnosis, we demonstrated widespread and early methylation changes pre-dating AML diagnosis. By integrating these epigenetic measurements with longitudinal deep, error-corrected genetic sequencing, we found tens of thousands of CpGs consistently associated with pre-leukaemic clonal expansions.

Our findings support a model wherein epimutations accumulate at a rate of approximately 0.1% – 1% per CpG per year across our targeted panel, 5-6 orders of magnitude higher than rates of somatic mutation ^*62*^. At these rates, methylation changes are thus widespread but remain highly heritable at a time-scale of decades. They therefore represent promising biomarkers that could substantially improve the sensitivity for the detection of pre-cancerous clones, complementing conventional somatic variants ^*81–84*^.

Our study supports two broad categories of epigenetic variation that accumulate in HSCs during ageing haematopoiesis. First, a class of functionally neutral fluctuating CpGs whose methylation states change stochastically ^*61*^. Our longitudinal data provides strong evidence supporting the idea that these fCpGs can be used for quantitative lineage tracing, building on previous work ^*60,61*^. These fCpGs may underpin an unbiased and cost-effective approach to tracing somatic evolution, with the advantage of potentially detecting cryptic clonal expansions. If such fCpGs could be phased using long reads, they could represent in vivo evolvable barcodes.

The second broad category of epigenetic variation we identify is likely to have a functional role in the early stages of AML development. We discovered hundreds of candidate epigenetic ‘driver’ regions that exhibit methylation changes specifically associated with AML and are recurrently shared among independent pre-AML cases. While it remains challenging to definitively determine whether these epigenetic alterations predispose cells to acquire genetic mutations, or conversely result from genetic events, two key findings strongly support a functional role for these regions in early disease. First, the driver DMRs display consistent methylation changes arising decades before clinical AML diagnosis, suggesting their involvement very early in disease progression. Second, these methylation alterations appear already fixed within the single HSC that initiates the pre-leukaemic clonal expansion, implying that these epigenetic changes are stable, heritable, and potentially influential in driving AML evolution. These findings are consistent with work showing that epigenetic changes acquired in cancer-associated genes during embryonic development predispose individuals to cancer later in life ^*85*^.

While many of the putative driver regions overlapped genes previously associated with AML, numerous novel regions were also discovered. One surprising finding was the large number of regions associated with neurodevelopmental genes (e.g., *ROBO1, SLIT2, GRIK2, PENK, ZIC4, ISLR2*). Previous work has suggested a role for neurodevelopmental genes in non-haematological cancers ^*86–88*^ and a role for cholinergic signalling in HSC maintenance ^*89*^. Our study adds to this evidence highlighting a potentially important role for these genes in AML initiation and progression, warranting further investigation.

Distinguishing true ‘driver’ regions from non-functional epigenetic variation is more challenging in the context of methylation than in the context of somatic genetic variation ^*90*^. Methylation patterns inherently show greater variability than genetic mutations, and the high intrinsic epimutation rate further complicates identifying functionally relevant recurrent events. While our stringent filtering approach aimed to minimise false positives, some regions identified as putative drivers could still represent sites prone to stochastic epigenetic alterations (‘epigenetic drift’). Future studies, particularly larger-scale studies in control populations with healthy clonal haematopoiesis, will be critical to rigorously validate these candidate driver regions.

Our findings establish a foundation for integrating methylation-based biomarkers with somatic genetic alterations to improve AML risk prediction and early detection. The identified epigenetic markers and pathways provide novel avenues for therapeutic discovery. Given the fundamental role of methylation in shaping cell identity across tissues, these approaches are likely applicable to other cancers.

## Methods

### BLUEPRINT samples and processing methodology

Whole genome bisulfite sequencing (WGBS) files were downloaded as unaligned BAM files through the EGA database (accession IDs EGAD00001002732, EGAD00001002419 and EGAD00001002333). Data from purified HSC samples from 8 healthy donors, purified blasts from 21 AML patients, and purified neutrophils, monocytes, CD4+ T-cells, CD8+ T-cells, B cells, and NK cells from 3 healthy donors was used. AML samples were from adult AML cases (age at diagnosis *∼*55) and offered a good representation of the mutational landscape of the disease. Unaligned BAMs were converted to FASTQ format using samtools ^*91*^ bam2fq. FASTQ files were mapped to the hg19 genomic annotation using Bismark ^*92*^, with the default settings and bowtie2. The resulting BAM files were deduplicated, using bismark, with the default settings applied. The bismark methylation extractor tool was used to generate methylation frequencies and coverage data for each CpG cytosine. The average depth was 4.2X across HSC samples, 18.5X across AML blast samples, and 1.5X across peripheral blood purified cell samples.

### DMR Identification Algorithm

The DMR identification algorithm was custom written using python. Firstly, depth filters were applied to the data. CpG cytosines covered to at least 3X in at least three AML donors and to at least 2X in at least two HSC donors were included in the analysis. *∼*28 million CpG cytosines passed this filter in both datasets and were used for discovery. For each cytosine passing the depth filter, a region was initiated. The difference in mean methylation between the two groups (up to n = 21 AML vs. up to n = 8 HSC, for BLUEPRINT data) in units of the standard deviation (*z*-score) was calculated for that root cytosine. This is given as:

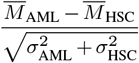

where:

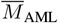= mean methylation in AML

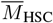= mean methylation in HSC

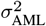= variance in AML

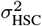= variance in HSC

The variance was upper-bounded, for both datasets, such that if the mean across that region was above 0.5, a “0” was added to the set of values used to calculate the variance, whereas if it was below 0.5, a “1” was added. The algorithm then extended one CpG cytosine at a time, in each direction separately, recalculating *z* for the region each time. Extension continued as long as the *z*-score increased. This process generated a candidate DMR around each CpG cytosine. A final *z*-score was then calculated for the entire extended region, and overlapping candidate DMRs with *z >* 3 were merged. For the merged regions, the final *z*-score and mean methylation difference were computed across all CpGs within the merged span.

### Validation ERRBS samples and processing

SRA files for enhanced reduced representation bisulfite sequencing (ERRBS) data for 119 AML sample from Glass *et al*. ^*49*^ (GEO GSE86952) were downloaded using the SRA toolkit (NCBI). These were transformed into FASTQ files using the fastq-dump command of the SRA toolkit. Further processing was identical to that described for BLUEPRINT samples, with the distinction that the deduplication step was omitted, as this is not recommended for ERRBS data. The depth was *∼* 60X per sample. For DMR validation, we considered only regions for which >33% of CpGs and >2 CpGs were covered in >40 donors. Region methylation was given as the mean across all CpGs in all samples. Regions were considered ‘validating’ if the methylation difference between the ERRBS AML samples and BLUEPRINT HSC samples was in the same direction as when comparing BLUEPRINT leukaemic blast samples to HSC samples (i.e. hypomethylated if previously hypomethylated, hypermethylated if previously hypermethylated).

### UKCTOCS samples

The UK Collaborative Trial of Ovarian Cancer Screening (UKCTOCS) (ISRCTN22488978; ClinicalTrials.gov NTC00058032) was a multi-centre randomised controlled trial set up to assess the impact of screening on ovarian cancer mortality ^*52*^. As part of this study, blood samples were collected annually for over 50,000 postmenopausal women over a period of up to 11 years: eligibility and trial details have been reported previously ^*93*^. UKCTOCS was reviewed and approved by the North West Multi-centre Research Ethics Committee (MREC) (REC reference 00/8/34) and participants provided written consent for use of their samples and data in ethically approved secondary studies. Our study, using UKCTOCS samples and data, was reviewed and approved by the South Central Berkshire B Research Ethics Committee (REC reference 18/SC/0481).

UKCTOCS blood samples were taken in Greiner gel separation tubes and transported at ambient temperature for processing. Due to an extended wait time between sample collection and centrifugation (median *∼*22.1 hours), leukocyte genomic DNA leaked into the serum ^*28*^, rendering samples suitable for genomic analyses. Partipants’ health outcomes were followed and, by 2018, 50 women with longitudinal sampling had incidentally developed AML (ICD-10 C92.0) (mean age 71). We obtained 1ml of serum from each of the yearly timepoints from these 50 women, as well as from 50 age-and timepoint-matched UKCTOCS participants who have remained cancer free. DNA was extracted by LGC Genomics, using an adapted KleargeneTM method with mag beads on KingFisherTM 96, and eluted in 10 mM Tris-Cl pH 8.0.

### WGBS library preparation and sequencing

WGBS libraries were prepared from a starting amount of 10ng per sample. Sonication was carried out using the M220 focused-ultrasonicator (covaris) and microtube-50 (covaris) tubes. Settings for sonication were set according to manufacturers’ specifications: peak incident power 75, duty factor 10, and cycles per burst 200. Each sample was sonicated for 70 seconds. The resulting products were transferred to a clean eppendorf tube, and concentrated using the zymo DNA clean and concentrator kit (D4013), to 10µL total volume. Bisulfite conversion was carried out using the EZ DNA Methylation-Lightning kit (zymo research D5030), according to manufacturers’ instructions. This was followed by adaptor tagging, extension, ligation, and indexing PCR (11 cycles) with the Accel-NGS® Methyl-Seq DNA Library Kit (swift biosciences 30024) and the Accel-NGS Methyl-Seq Unique Dual Indexing Kit (swift biosciences 390384), according to manufacturers’ instructions. Libraries were sequenced on an Illumina NovaSeq S4, with paired end reads (2×150bp) at *∼* 250 million reads per sample, with a 25% phiX spike-in (CRUK Cambridge Institute Genomics Core Facility). The samples were demultiplexed by the core, following the sequencing run. The resulting FASTQ files were first trimmed to remove adaptors and 10bp from each end, using TrimGalore ^*94*^, then processed as described for BLUEPRINT datasets, but with default settings for paired-end sequencing.

### UKCTOCS samples cell deconvolution

WGBS libraries were prepared for 96 UKCTOCS samples from 46 donors (23 cases and 23 controls). To estimate the proportions of blood cell types present in the samples (B-cells, NK-cells, CD4T and CD8T-cells, monocytes, neutrophils and eosinophils), we applied the R package EpiDISH using the reference dataset “centDHS-bloodDMC.m” within the package ^*95*^. As this approach leverages methylation frequency across 333 CpGs, we considered all samples for which at least 286 CpGs were covered (88 samples, average depth *∼*7X).

### Enrichment for regulatory features

For analysis of genomic context and feature enrichment, all information was obtained from the UCSC Genome Browser ^*96*^, for alignment hg19. For the enrichment analysis, in order to be classified as having a signal for H3K27ac or for the GeneHancer the score at the site had to be in the 90th percentile for the feature in question.

### Gene ontology enrichment analysis

DMRs were annotated as pertaining to a specific gene where there was overlap with the gene coordinates (including the promoter as the 1Kbp region upstream of the start site), obtained from the UCSC Genome Browser ^*96*^ NCBI Refseq track. Gene ontology enrichment analysis was carried out using the python package GOATOOLS ^*97*^. A bonferroni-corrected p-value*<* 0.01 was set as the threshold to define the functional enrichment significance.

### Literature review of genes associated to DMRs

For each gene linked to a DMR, a pubmed search was conducted with the terms: “<gene> methylation acute myeloid leukaemia” and “<gene> acute myeloid leukaemia". This was automated using Python, where search results were retrieved via the Entrez API (to a maximum of ten results). The articles returned were manually checked, to ascertain whether there was previous evidence for methylation or expression changes in AML.

### Developing a targeted panel of AML-relevant regions

An overview of the rationale and filtering details for each type of region included in the panel is shown in Extended Data Fig. 7). Five types of regions were included. (i) Regions defined from leukaemic blasts/HSCs and validated in peripheral blood: All hypo-and hypermethylated DMRs of *z >* 3 identified from leukaemic blast and HSC datasets were considered if the methylation difference between the peripheral blood pre-AML and control groups was in the expected direction. Given the confidence in leukaemic blast-defined DMRs, especially the hypermethylated group, we assigned just under 50% of all panel probes to leukaemic blast-defined DMRs, with an *∼*25/75% split for hypo- and hyper-methylated DMRs, respectively. As a high number of regions passed the thresholds set, DMRs were ranked by the *z*-score of the DMR in peripheral blood (based on WGBS data from 23 pre-AML cases compared to 23 age-matched controls). Regions were included in the panel until the appropriate quota of probes were used. (ii) Regions defined from peripheral blood samples: DMRs were also discovered by applying the DMR-identification algorithm described above to WGBS data for 23 pre-AML cases compared to 23 age-matched controls. We allocated just under 25% of probes to peripheral blood-defined regions. (iii) High entropy regions: Disorder was calculated for each 4-CpG locus using Shannon entropy ^*98*^ in 6 leukaemic blast donors. Loci with high ME (*>*0.6/1) in these donors which validated in the remaining leukaemic blast donors were merged. Just under 25% of probes covered regions defined by ME. (iv) AML gene promoters: probes were included which covered the promoter regions of 35 genes linked to AML and CH (Supplementary Table 1). The promoter regions were considered to be the 1000bp upstream of the start codon. Where a gene had alternative promoters, all were included. Finally, clock CpGs from the skin and blood clock ^*56*^ were covered, accounting for *∼* 2% of the targeted panel. Regions which overlapped were merged. Final panel design was carried out by Twist biosciences, with any targets in repetitive regions removed.

### Targeted DNA methylation library preparation and sequencing

Sequencing libraries were prepared from 30ng of input DNA. Sonication was carried out as described for the WGBS library preparation. Following sonication, whole-genome libraries were prepared using the NEBNext® EM-seq kit (NEB E7120), according to manufacturer’s instructions, with 10 cycles of PCR in the final amplification step. Targeted enrichment was carried out using the Twist Bioscience Targeted Methylation Sequencing workflow, according to manufacturer’s instructions, using a custom-designed set of probes, as described above. Paired end sequencing (2×150bp) was carried out by the genomics core at the CRUK Cambridge institute. Samples were sequenced on a NovaSeq S4 flow cell, at *∼* 75 million reads per sample, with a 10% phiX spike-in. Samples were demultiplexed by the genomics core.

FASTQ reads were adaptor trimmed, followed by a 5’ quality trim of 4bp, and a 3’ quality trim of 1bp, using TrimGalore ^*94*^. FASTQ reads were aligned to the hg19 reference genome annotation using Bismark ^*92*^, with the default settings for paired-end sequencing and bowtie2. Following this, the resulting BAM files were deduplicated using bismark, with the default settings for paired-end sequencing applied. Samples with CHH meth *>* 3% or average depth *<* 200X were excluded from the analysis: 475 samples passed these filters. The average depth across these samples was *∼* 773X per sample (range 204-1687).

### DMR trajectories

For each sample, the average methylation at a CpG was calculated using all reads from both strands. The methylation across the region in each sample was then calculated as the mean of all CpGs in that region (including only CpGs at depth >100X). In all regions where at least 50 control samples were covered at high depth (*n* = 5,120 regions), we calculated the expected mean methylation and the standard deviation of the mean across control samples using region-level methylation frequencies. For each case sample, a normalised methylation difference was then computed as:

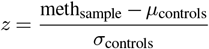

where:

meth_sample_ = mean methylation frequency for the region in the sample

*µ*_controls_ = mean of region-level methylation frequencies across all control samples

*σ*_controls_ = standard deviation of region-level methylation frequencies across all control samples

For control samples, the normalised difference was calculated in the same manner, except that any samples from the control in question were excluded from the calculation of the control mean and standard deviation.

### Somatic sequencing

Integrative duplex error-corrected DNA sequencing (‘TETRIS-seq’) was carried out as previously described ^*59*^. This approach incorporates two targeted panels (Twist Biosciences). One is a small (*∼*58kb) panel specifically for SNVs and indels, targeting commonly mutated exons within 34 genes commonly mutated in clonal haematopoiesis and AML: ASXL1, BCOR, BCORL1, CBL, CEBPA, CHEK2, CSF3R, DDX41, DNMT3A, EZH2, FLT3, GATA2, GNAS, GNB1, IDH1, IDH2, JAK2, KIT, KRAS, MPL, NPM1, NRAS, PPM1D, PTPN11, RAD21, RUNX1, SF3B1, SRSF2, STAG2, TET2, TP53, U2AF1, WT1, ZRSR2. The other is a large (*∼*1.6 MB) panel targeting a total of 10,326 common SNPs (minor allele frequency > 0.01) spaced every *∼*280 kb across the genome for the detection of mosaic copy number alterations (mCAs) and common breakpoint regions for detection of 8 common AML chromosomal rearrangements: t(6;9) DEK::NUP214, t(8;21) RUNX1::RUNX1T1, t(9;11) KMT2A::MLLT3, t(9;22) BCR::ABL, t(15;17) PML::RARA, t(16;16) CBFB::MYH11, inv(16) CBFB::MYH11 and KMT2A partial tandem duplication. A custom in-silico de-noising algorithm was applied to the resulting data, enabling detection of single nucleotide variants (SNVs) and indels at variant allele frequencies (VAFs) *≥* 0.1%, and of chromosomal gains, losses and copy-neutral loss of heterozygosity (CN-LOH) events down to 0.1% cell fraction.

### Establishing associations between somatic variants and CpG methylation through time

To determine whether the methylation levels of individual CpGs presented a significant association to the variant cell fraction (VCF) trajectory of specific somatic variants, we considered competing hypotheses. Under the null hypothesis, methylation levels are assumed to remain constant (equal to their mean) over time. Under the alternative hypothesis, CpG methylation frequency over time is driven by the somatic clone expansion, assuming a constant methylation level for that somatic clone (meth_*mut*_) and for the background (meth_*W T*_). We employed a constrained least-squares optimisation approach to determine the best-fit methylation levels for both the somatic driver clone and the background, using the optimize.lsq_linear function from the python package scipy. To evaluate the improvement in fit provided by this approach over the null hypothesis, we considered the residual sum of squares (computed as the sum of squared differences between the observed and expected methylation levels) for each model, and calculated an F-statistic as:

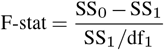

where:

SS_0_ = residual sum of squares under the null hypothesis (methylation mean static over time)

SS_1_ = residual sum of squares under the alternative hypothesis

df_1_ = degrees of freedom, calculated as the number of data points minus two parameters inferred under the alternative hypothesis

This was applied to every variant in each donor, provided a change in VCF of at least 3% through time, for each CpG. To determine the F-statistic threshold above which a CpG was considered associated to each variant, we permuted the observed methylation once for each of the ∼150, 000 CpGs, and recalculated the best-fit methylation levels and F-statistic using the permuted values. The F-statistic threshold was taken as the lowest value at which the proportion of CpGs above that value under permuted conditions relative to the number of CpGs above that value considering the true methylation values was under the desired false-discovery rate. Where this was achieved at any threshold, 17 variants across 10 donors passed an FDR threshold of 5%, and variants in a further two donors passed an FDR threshold of 10% (Fig. 3). In certain donors (C92_002 and C92_007) variants are present on the same subclone at similar VCFs through time, so it is not possible to distinguish CpGs associations between them. In these cases, we considered only the associations with the most recently acquired variant, i.e. the one characterising the subclone with all variants. To compare the predictions of the alternative hypothesis to the ground truth (Fig. 3e-q), we plotted the expected VCF frequency, given as:

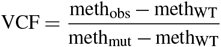

where:

meth_obs_ = observed methylation through time for a specific donor and

CpG meth_mut_ = best-fit methylation level of the driver clone

meth_WT_ = best-fit methylation level of the background

### Estimation of epimutation rate

To estimate the epimutation rate across our panel, we considered only donors in which the wild-type background was dominant at the first timepoint, and a somatic variant increased in frequency to *>* 20% VCF by the final timepoint. Based on the trajectories of the somatic variants within a donor, fitness and mutation establishment time can be inferred for each clone detected (see Watson *et. al*. ^*59*^: in press). For each donor passing the criteria set, we considered the clone for which the somatic variant associated to is the most recently acquired. The epimutation rate is then given as:

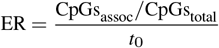

where:

CpGs_assoc_ = number of CpGs statistically associated with the variant (as described above)

CpGs_total_ = number of CpGs tested (i.e., all panel CpGs passing a coverage filter of 100X, *∼*150,000)

*t*_0_ = establishment time

The error on this measurement is calculated using propagation of uncertainty:

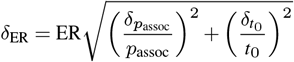

where:

*p*_assoc_ = proportion of CpGs associated with the clone (CpGs_assoc_*/*CpGs_total_)

*δ*_*p*assoc_ = uncertainty in the proportion of associated CpGs, calculated as:

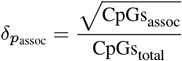

and 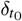 = uncertainty in the establishment time, given by:

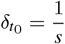

where *s* represents the clone fitness.

### Calculation of epimutation burden

To identify CpGs significantly altered relative to the expected frequency levels in each sample, we considered only CpGs covered at *>* 100X in at least 50 case samples and at least 50 control samples (∼150, 000 CpGs). In each individual sample, we considered only CpGs covered at *>* 100X in that sample. For case samples, the expected mean methylation and standard deviation of the mean were calculated using the frequencies from all control samples. The *z*-score for each CpG was then calculated as:

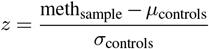

where:

meth_sample_ = methylation frequency of the CpG in the sample

*µ*_controls_ = mean methylation frequency of the CpG across all control samples

*σ*_controls_ = standard deviation of the mean methylation frequency of the CpG across all control samples

For control samples, the z-score was calculated in an identical manner, but any samples from the control in question were not included in the calculation of mean and standard deviation. A CpG was considered altered in a sample if the z-score was above 6. The epimutation burden in each sample is the proportion of highly-covered CpGs which are altered.

### Defining AML-specific DMRs

For each sample and region, we considered how many CpGs were altered (z-score >6, see previous section). A region was considered differentially methylated in a sample if >10% of CpGs and >2 CpGs were altered (NB: this excluded short regions by default), and all CpGs altered were altered in the same direction (i.e. all hyperor all hypomethylated). For a region to be considered AML-specific, no control samples could meet this criteria, while samples from more than five separate pre-AML donors did so. The normalised difference in AML-specific DMRs (Fig. 4c-g) was given as the mean z-score across all CpGs in the region.

### Discovery of putative fCpG loci

To identify potential fCpG sites covered by our targeted enrichment panel, we considered all CpG positions, independent of strand orientation, with *≥*100X coverage in all donor libraries, resulting in 405,192 assessable CpGs. Next, we used genetic sequencing results generated by Watson et al. ^*59*^ to identify the case libraries with high, somatic variant allele fractions (VAFs) and control libraries with no detectable somatic mutations, indicating a clonal expansion and no clonal expansion, respectively. We selected four case libraries with the highest somatic VAFs at the final blood draw – referred to as ‘discovery cases’ (C92_022_s7, C92_033_s7, C92_034_s7, and C92_062_s3; see red asterisks in Fig.5d) – and randomly chose four of the 13 control libraries with no detectable somatic mutations, designated as “discovery controls” (CNTRL_002_s4, CNTRL_003_s2, CNTRL_165_s1, and CNTRL_171_s1; see blue asterisks in Fig.5d). We then selected CpG sites with mean methylation fractions between 0.4 and 0.6 in *≥*3 of the 4 discovery control libraries, resulting in 21,643 CpG sites. To generate our final set of putative fCpGs, we further filtered those sites such that each CpG site had a mean methylation fraction of *≤*0.1 in *≥*1 of 4 discovery case libraries, *≥*0.8 in *≥*1 of 4 discovery case libraries, and between 0.3 and 0.6 in the remaining *≤*2 discovery case libraries. Taken together, these constraints resulted in 1,045 putative fCpG loci (see Supplementary Table 2).

### Assessment of putative fCpGs in non-discovery cases and controls

We assessed the performance of our 1,045 putative fCpG sites by comparing the methylation distributions of these sites in libraries with and without known clonal expansions. Samples were omitted if they had *≤*900 fCpGs covered at *≥*100X depth. We calculated the variance of the distribution of fCpG methylation fractions (abbreviated as “fMV”) for each library as an approximate measurement of that distribution’s “W-ness” and compared it the cell-fraction of largest clone in that library. The relationship between the fMV and the clonal cell fraction (CCF) of largest clone was predicted to be quadratic when W-shaped distributions were modelled as a mixture of three Gaussian distributions. To assess the relationship between fMV and clonal burden in C92_002 blood draws, we fit a second-order polynomial regression (with the linear component constrained to zero) to the fMV and largest CCF (*R*^2^=0.997, see Fig. 5c).

### fCpG data-fitting and identification of donors with likely missing drivers or miscalled germline mutations

Based on our observations of high fMV in some pre-AML samples with no detectable somatic mutation we sought to develop a principled method for identifying outlier samples resulting from (i) insufficient detection of somatic clonal markers (i.e., missing driver mutations), (ii) germline mutations incorrectly called as somatic (i.e., miscalled germline), or (iii) very early somatic sweeps (>20-30 years prior to sampling) in which synchronized (i.e., clonal) fCpGs have had ample time to epimutate and become desynchronized. To this end, we assessed the quantitative relationship between fMV and clonal burden in donor blood samples using the combined data from three pre-AML cases that had mutation-associated clonal sweeps and increases in fMV during the sampling period (C92_002, C92_008, and C92_041). For each library in the combined data, we modeled this relationship using a second-order polynomial (as above, *R*^2^=0.974) and generated 95% confidence intervals (CIs, Extended Data Fig. 14a). To extend CIs to *∼*0% and 100% CCF, we assumed residuals for these values were the same as the nearest existing points. Next, we expanded the range of the CIs for a given CCF value in the model to include *∼*85% of our data (11-fold) and defined libraries as outliers if they were outside of these thresholds (Extended Data Fig. 14b). Outliers above this region are predicted to have a clonal expansion with an undetected somatic driver mutation, and those below it are suspected germline mutations that were miscalled as “somatic.” We used this polynomial model to predict the clonal burden (largest CCF) for each library based in its fMV (Extended Data Fig. 14c) where large deviations from y=x were assumed to be a result of missing drivers, miscalled germline, or very early somatic sweeps.

## ACKNOWLEDGMENTS

We acknowledge Dan Landau, Inigo Matrincorena, Jyoti Nangalia, George Vassiliou, Wolf Reik, Charlie Massie, Tom Mitchell and all members of the Blundell lab for helpful comments on this work. J.R.B and A.V.A.F are supported by the Early Cancer Institute, the CRUK Cambridge Centre and the NIHR Biomedical Research Centre. J.R.B. is supported by a UKRI Future Leaders Fellowship (MR/S031782/1). A.V.A.F and C.B are both supported by pathway awards from the International Alliance for Cancer Early Detection, an alliance between Cancer Research UK (C14478/A29329), Canary Center at Stanford University, the University of Cambridge, OHSU Knight Cancer Institute, University College London and the University of Manchester. C.J.W. was supported by a CRUK Clinical Research Fellowship and is currently supported by a Wellcome Early Career Award (226929/Z/23/Z). U.M., S.A. and A.GM. are supported by the NIHR UCL Hospitals Biomedical Research Centre and by Medical Research Council Clinical Trials Unit at UCL core funding (MR_UU_12023). TG and CG acknowledge funding from Cancer Research UK (DRCNPG-May21_100001 to TG, and EDDPMA-May23/100059 to TG and CG). We thank the UKCTOCS participants, without whom this study would not have been possible, and everyone involved in the conduct and oversight of UKCTOCS. UKCTOCS was funded by Medical Research Council (G9901012 and G0801228), Cancer Research UK (C1479/A2884), and the UK Department of Health, with additional support from The Eve Appeal. The long-term follow-up UKCTOCS was supported by National Institute for Health Research (NIHR HTA grant 16/46/01), Cancer Research UK, and The Eve Appeal. None of the funders had a role in data analysis, decision to publish or preparation of the manuscript.

## SUPPLEMENTARY MATERIALS

**Extended Data Figure 1.**
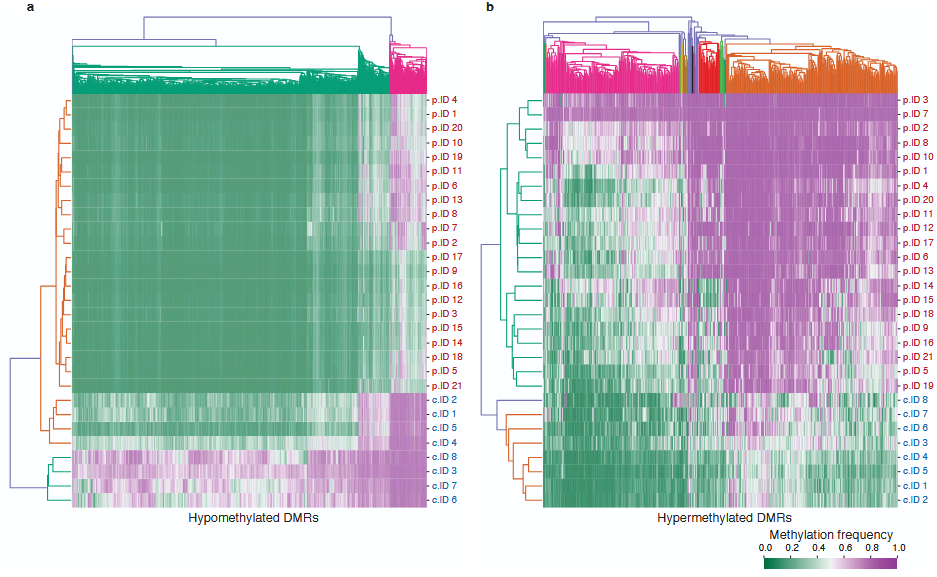
Methylation of identified DMRs across purified blast and HSC samples. Clustermaps and dendrograms showing hierarchical clustering of donors (rows) and DMRs (columns) for hypomethylated DMRs (**A**) and hypermethylated DMRs (**B**) of Z>6. AML samples are labelled in red and HSC samples in blue (right hand side).

**Extended Data Figure 2.**
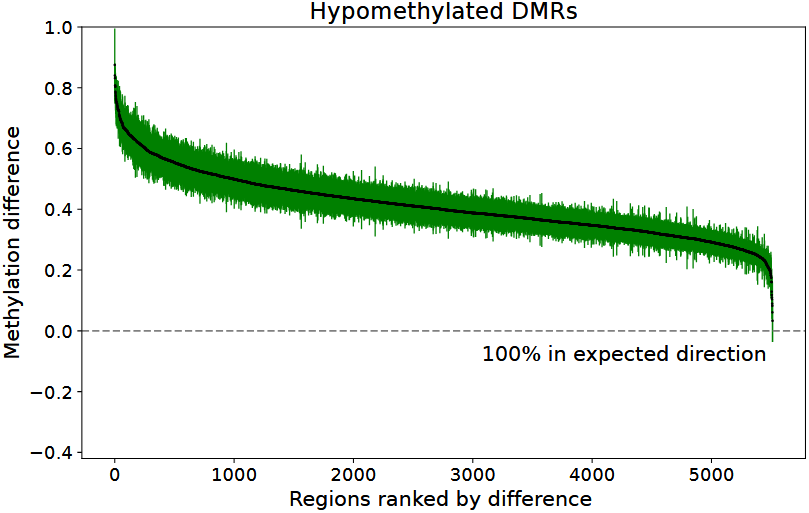
Validation of hypomethylated DMRs in an independent AML cohort. Mean methylation difference (black points) between an independent cohort of AML samples from Glass *et al*. ^*49*^ and BLUEPRINT ^*50*^ HSC samples, for all hypomethylated regions of z>6 identified from our discovery. Coloured bars show standard deviation and direction of difference. Results are representative of all DMRs, with 99.6% in expected direction at z>3.

**Extended Data Figure 3.**
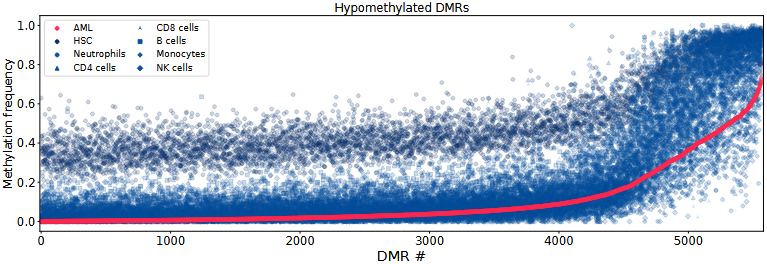
Hypomethylated DMR methylation levels in purified peripheral blood cells. Mean methylation level of each DMR of Z>6 across AML blasts (red), HSCs, and multiple peripheral blood cell types (blues). Results are representative of all DMRs (z>3).

**Extended Data Figure 4.**
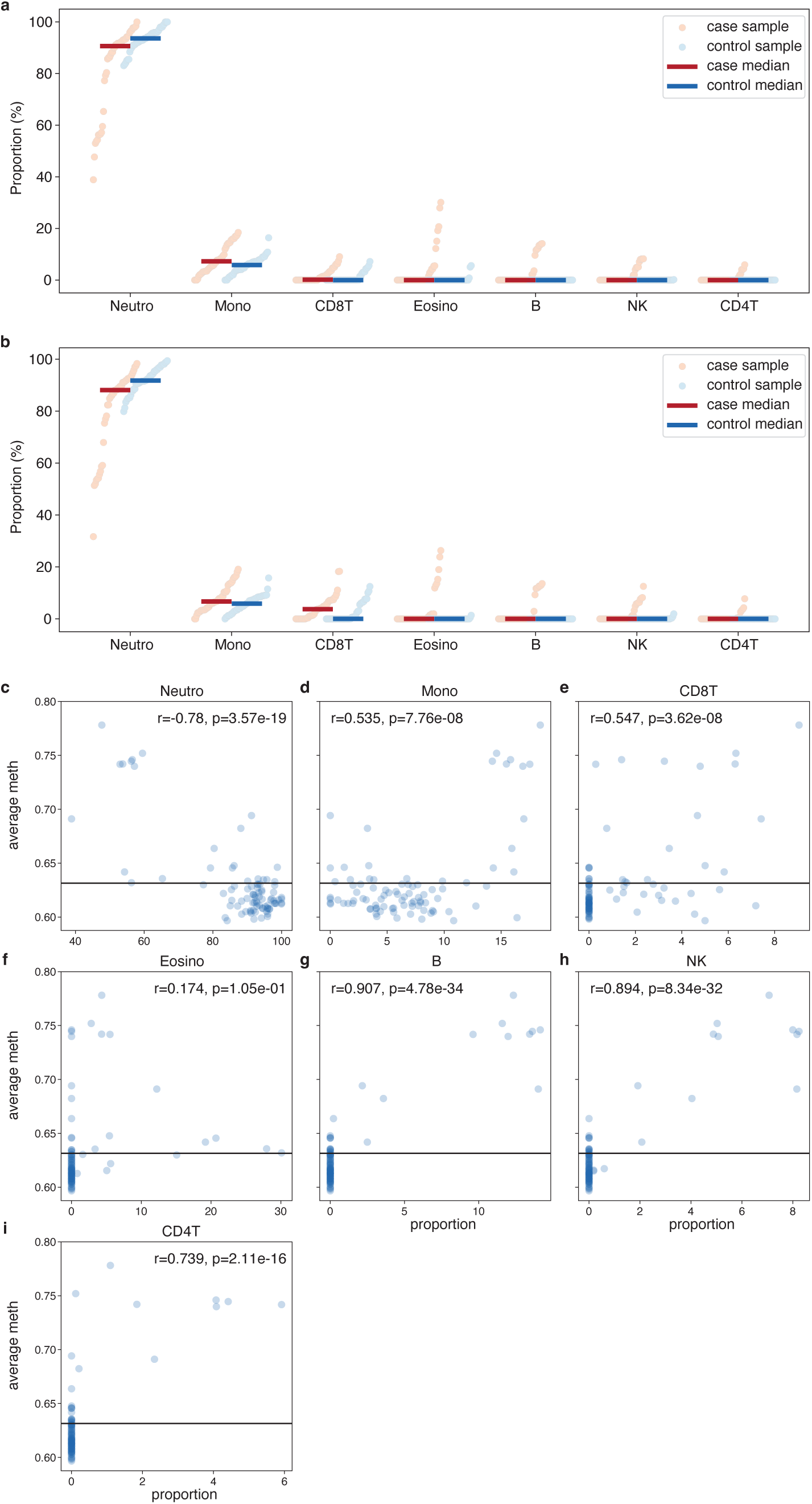
UKCTOCS cell deconvolution a and b. Estimates of proportions of each peripheral blood cell type in each sample (points), using the EpiDISH ^*95*^ package with method CBS (a) and RPC (b). Bars represent the median for each cell type across pre-AML samples (red) and controls (blue). **d-i** Relationship between cell proportion estimated and mean methylation in the sample, showing that the case outliers observed correspond to hypermethylated samples: i.e., this represents a disease signal rather than a true change in cell proportions. Black horizontal lines represent mean methylation levels in these samples for the CpGs considered. Pearson’s r and associated p-value shown for each cell type.

**Extended Data Figure 5.**
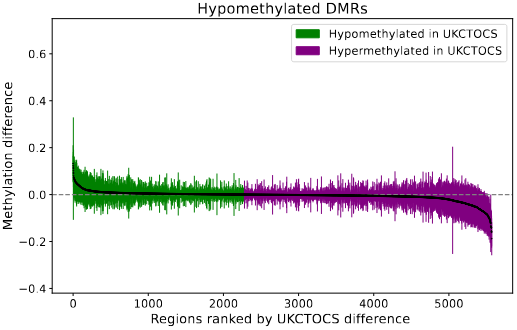
Validation of hypomethylated DMRs in peripheral blood pre-AML samples and controls. Mean methylation difference between pre-AML cases and matched controls (black points), for all hypermethylated regions of z>6. Coloured bars show standard deviation and direction of difference. Results are representative of all DMRs: 41% of DMRs in expected direction at z>6, 49% at z>3.

**Extended Data Figure 6.**
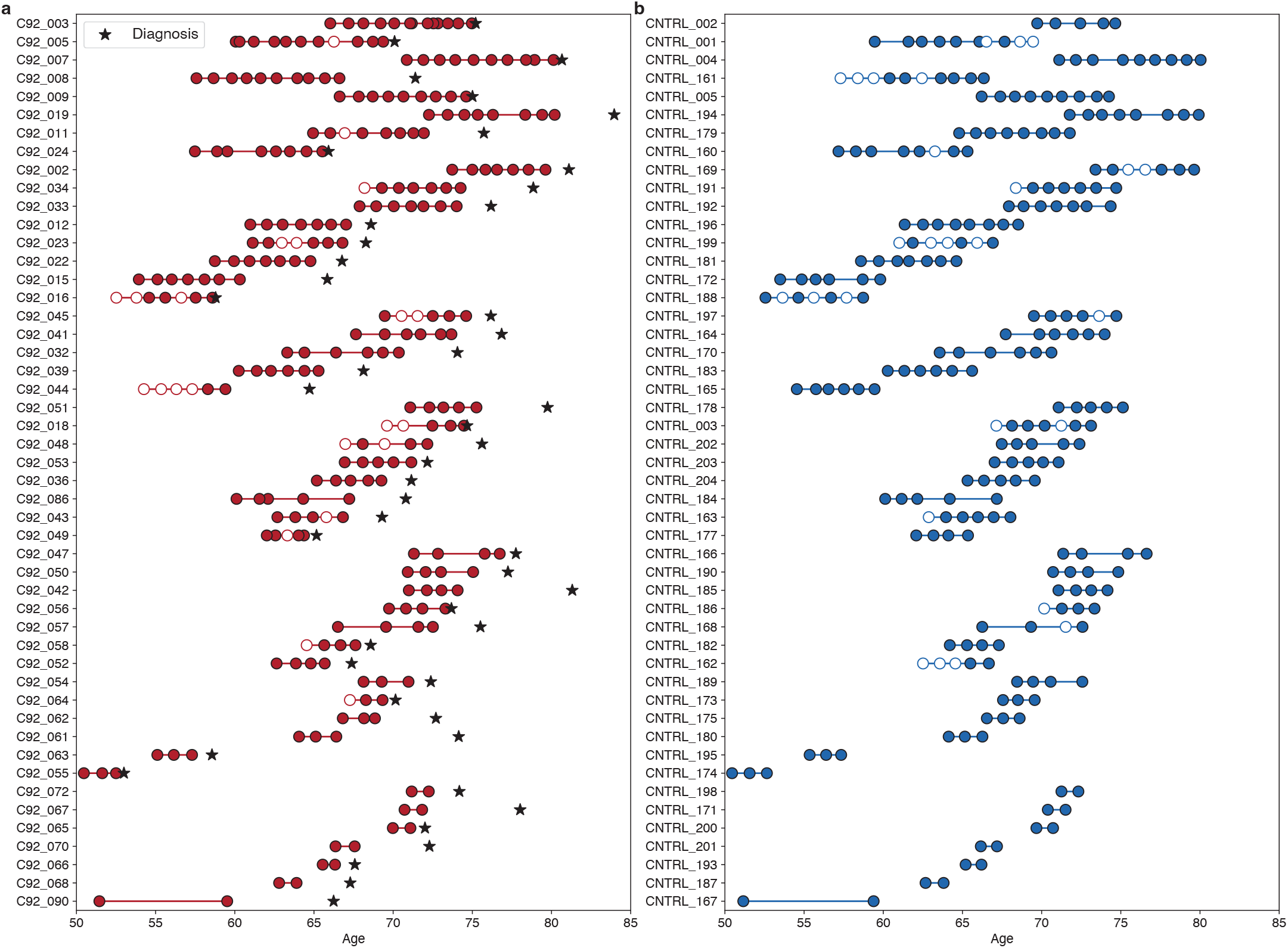
UKCTOCS samples. Pre-AML (**a**) and matched control (**b**) samples from the UKCTOCS study sequenced for this work. Each donor is shown as a line with circles representing individual samples. Filled samples passed depth and conversion filters (Methods).

**Extended Data Figure 7.**
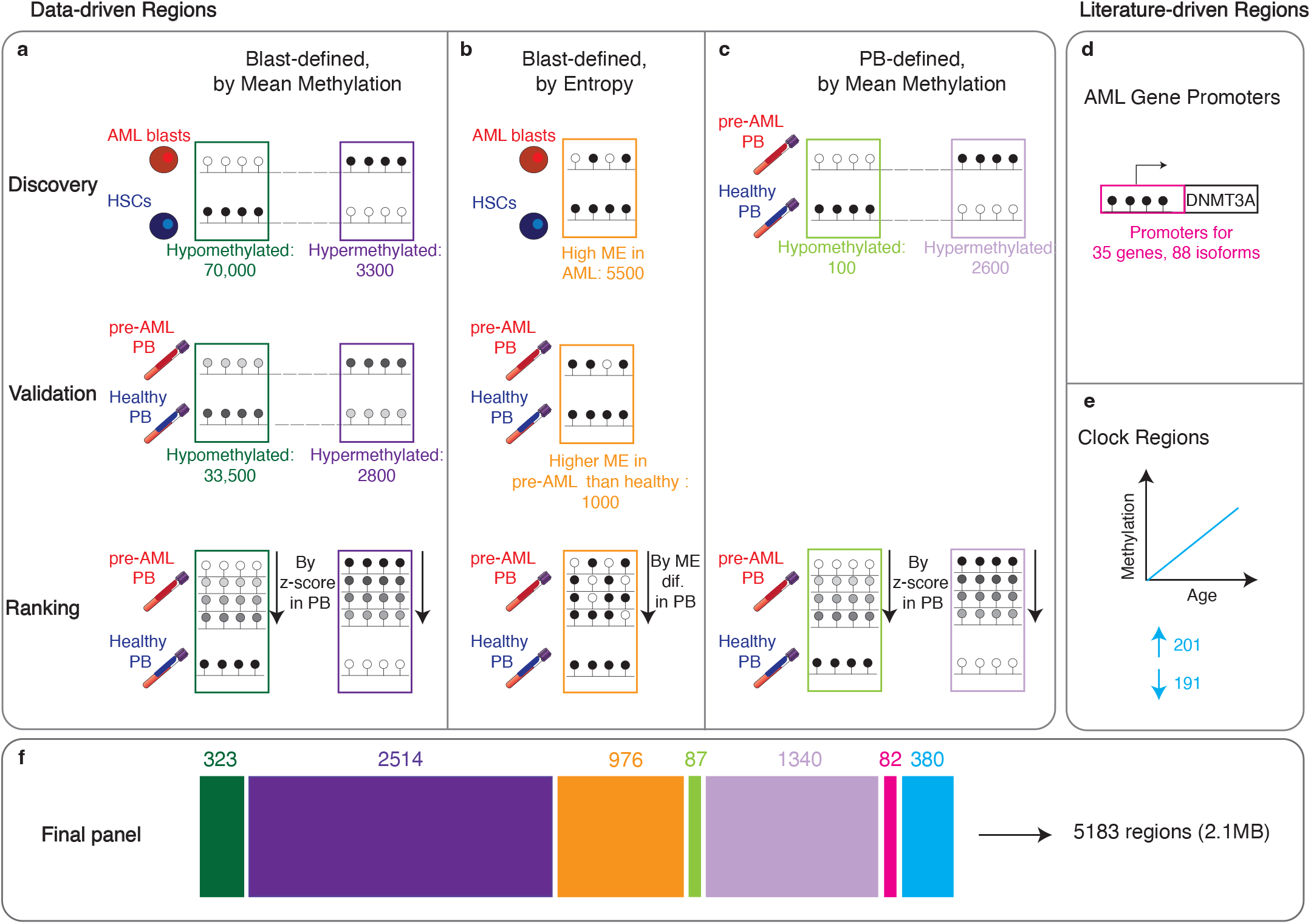
Probe panel design. **a** Regions identified as having a significant mean difference between leukaemic blasts and HSCs were filtered based on methylation difference in pre-AML vs. control PB, and ranked by the significance score of the difference. **b** Regions identified as showing high entropy in leukaemic blasts and low entropy in HSCs were filtered and ranked based on ME difference between pre-AML vs. control PB. **c** Regions identified as having a significant mean difference between pre-AML and control PB were ranked by the significance of the difference. Cut-offs for regions included in panel were based on probe quotas (main text). Lollipops represent CpG, with shade corresponding to methylation: white=unmethylated, black=fully methylated. **d** Promoter regions for known AML driver genes and **e** clock regions, for which the methylation level changes as a function of age, were added to the panel. **f** Final proportions and numbers of regions of each type, post Twist filtering and probe design. Colour legend: Dark green= blast-defined,hypomethylated. Dark purple= blast-defined, hypermethylated. Yellow= blast defined, entropic. Light green= PB-defined, hypomethylated. Light purple= PB-defined, hypermethylated. Pink= AML driver gene promoters. Blue= clock regions.

**Extended Data Figure 8.**
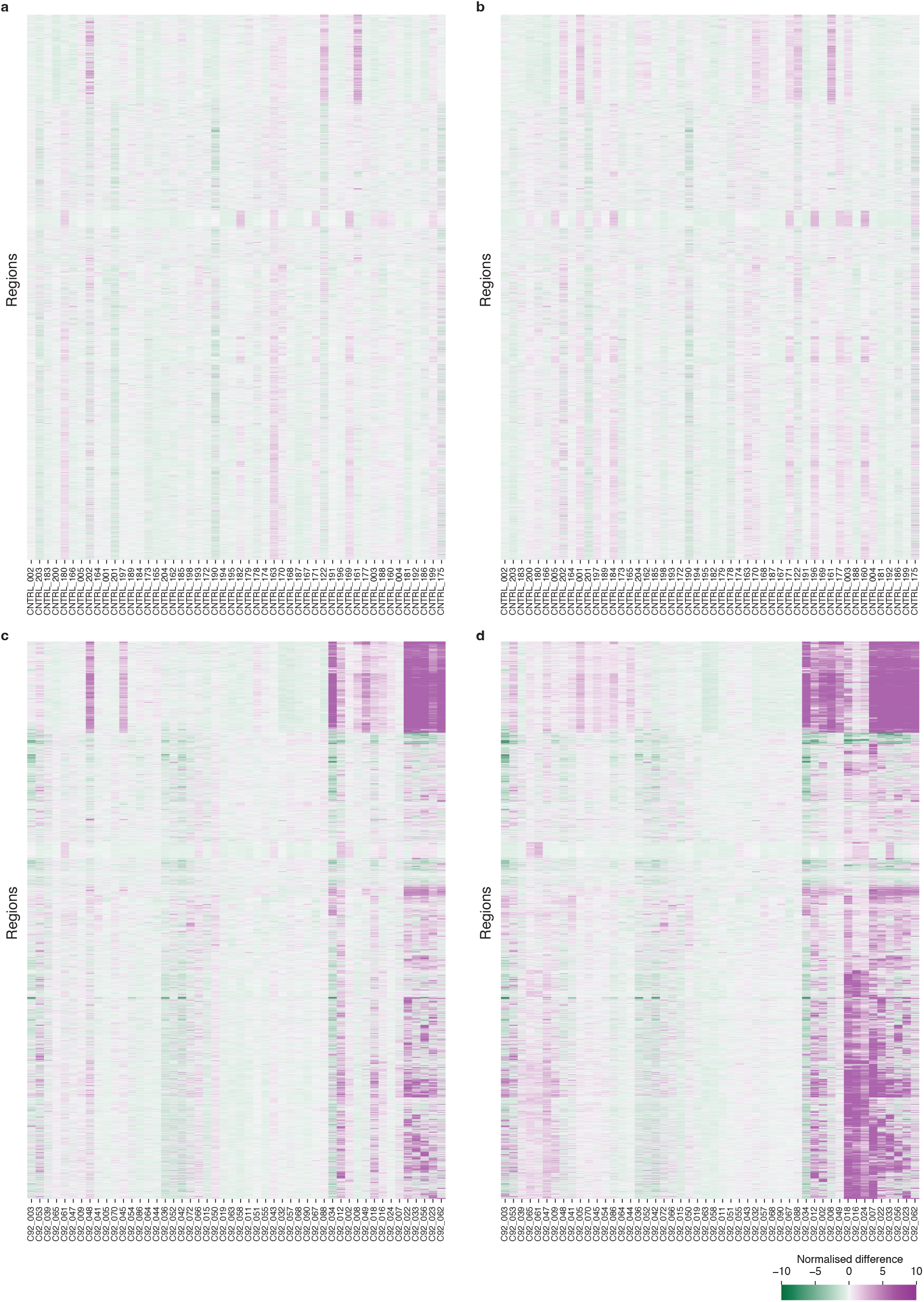
Region-level normalised difference from controls. Heatmaps showing the normalised difference (in units of the standard deviation) of the average methylation across each region between the sample in question and all controls (or all controls excluding the one tested, for control samples). **a** first timepoint for each control, **b** last timepoint for each control, **c** first timepoint for each case, **d** last timepoint for each case. All panel regions covered at the first and last timepoint for every donor are included (n=4825). Region and donor order is maintained and based on clustering at final timepoint, using the seaborn clustermap function with method=’ward’. Controls are ordered by their matched case.

**Extended Data Figure 9.**
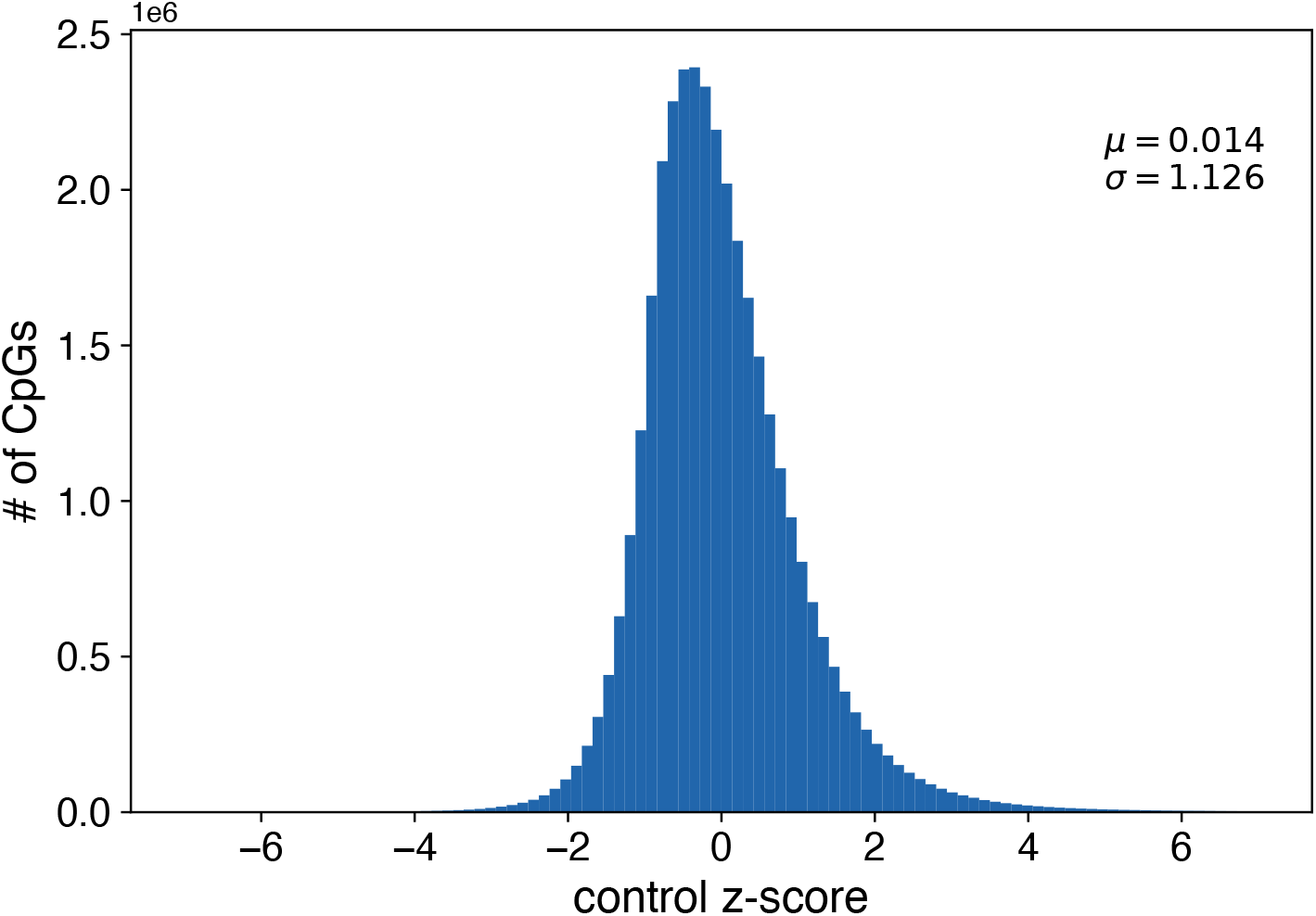
z-scores for all targeted panel CpGs across all control samples. The z-score for each CpG in each control sample is given as the difference between the methylation frequency of that CpG in that sample to the mean of all control samples excluding those from the control in question, in units of the standard deviation of the methylation frequency across all control samples considered. Mean and standard deviation of the z-score across all CpGs and all control samples are shown on plot.

**Extended Data Figure 10.**
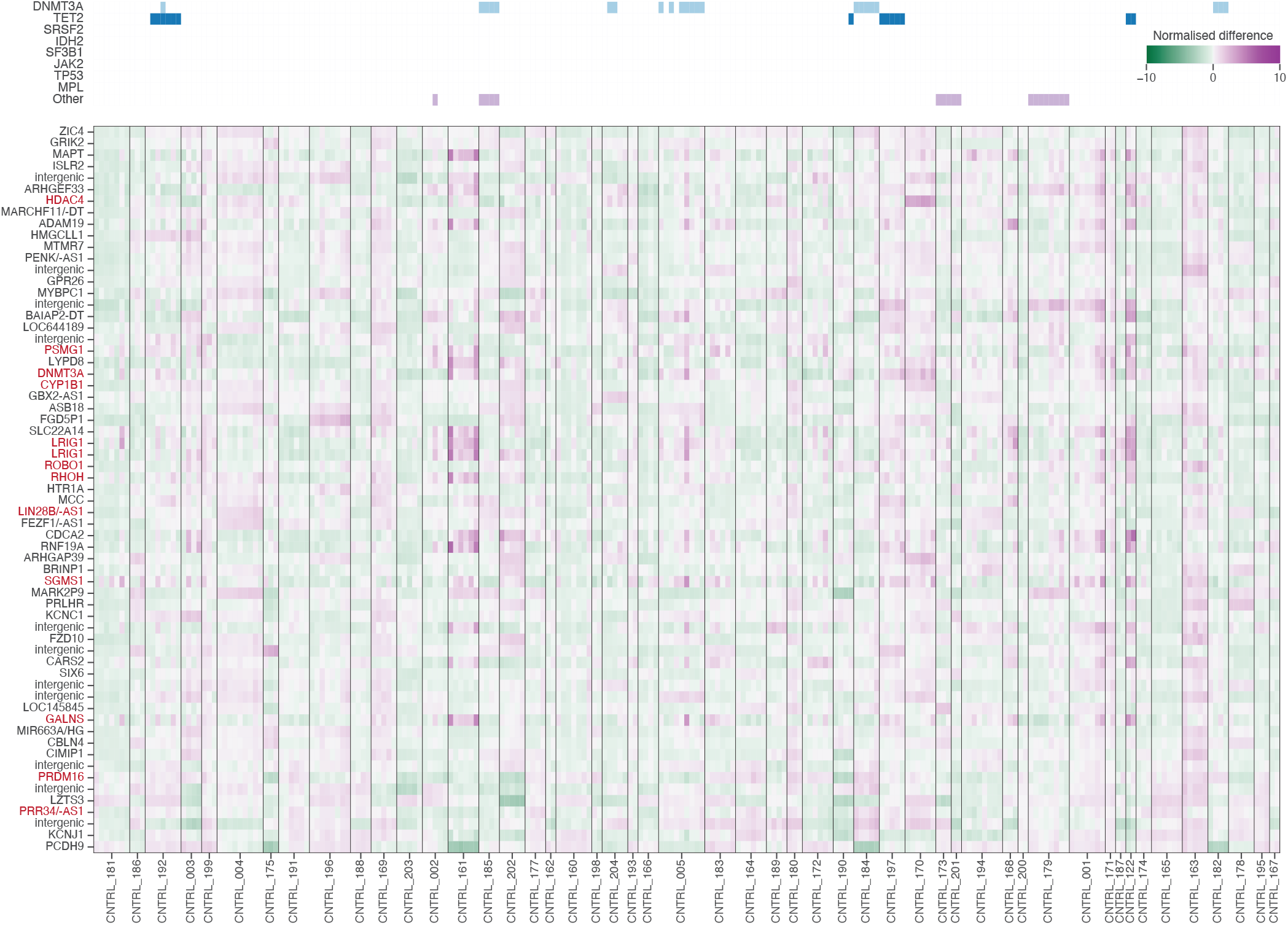
Control z-scores in highly recurrent AML-specific DMRs. Heatmap showing region *z*-score (mean of all CpGs in region) across regions (rows) and control samples (columns) for regions highlighted in Fig. 4c. Controls are ordered by their matched cases, with black lines separating donors. Column colours indicate presence of a somatic variant at above 5% VAF. Genes highlighted by red text have been previously linked to AML biology. Where labelled as <gene>/-AS1, both the gene and its antisense transcript intersect the region.

**Extended Data Figure 11.**
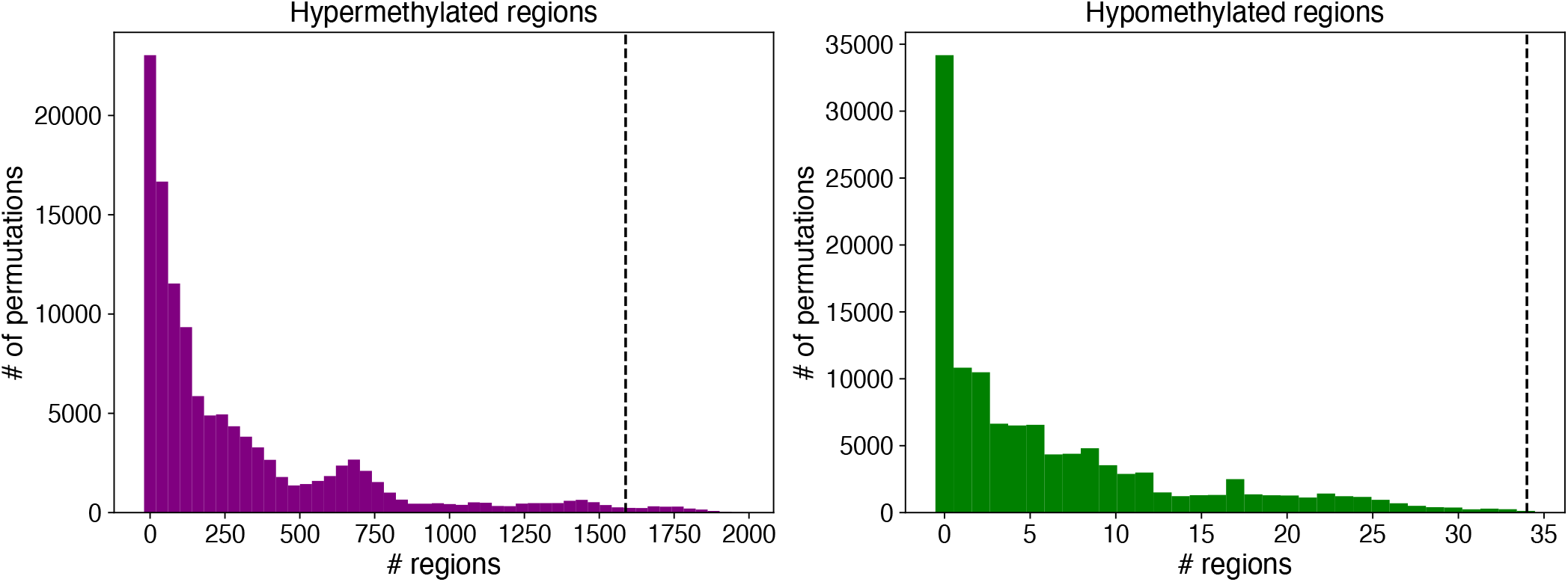
Numbers of “driver” regions when cases and controls are permuted. Considering only donors with an expansion (VAF*>* 0.4 or fMV*>* 0.05, 24 cases and 5 controls) these were permuted into all possible sets of 24 vs. 5. Plots show distribution of number of regions where no “controls” have methylation levels significantly changed from background level (methods) but over five “cases” do, across all possible permutations (118,775). **a** shows hypermethylated and **b** hypomethylated regions. Dotted line shows the number of regions passing filters when we consider the true case/control status across all 100 individuals.

**Extended Data Figure 12.**
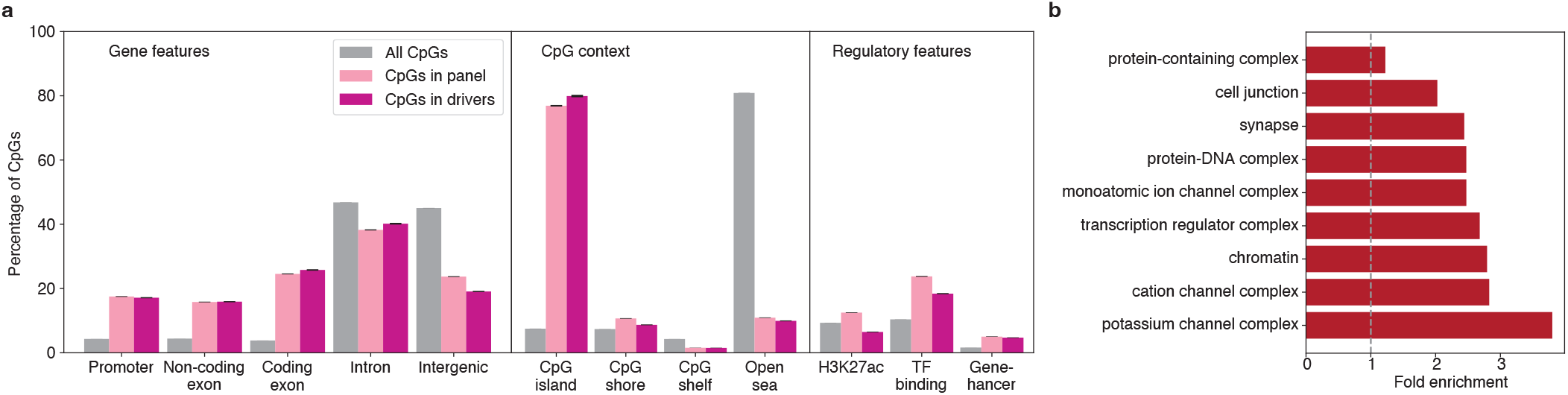
Feature enrichment in highly recurrent AML-specific DMRs. **a** Fraction of all CpGs (grey), CpGs within panel (light pink), and CpGs in highly recurrent “driver” regions (dark pink) for the labelled feature. **b** Fold enrichment for GO terms found to be significantly enriched for the set of genes overlapping driver regions (relative to all genes).

**Extended Data Figure 13.**
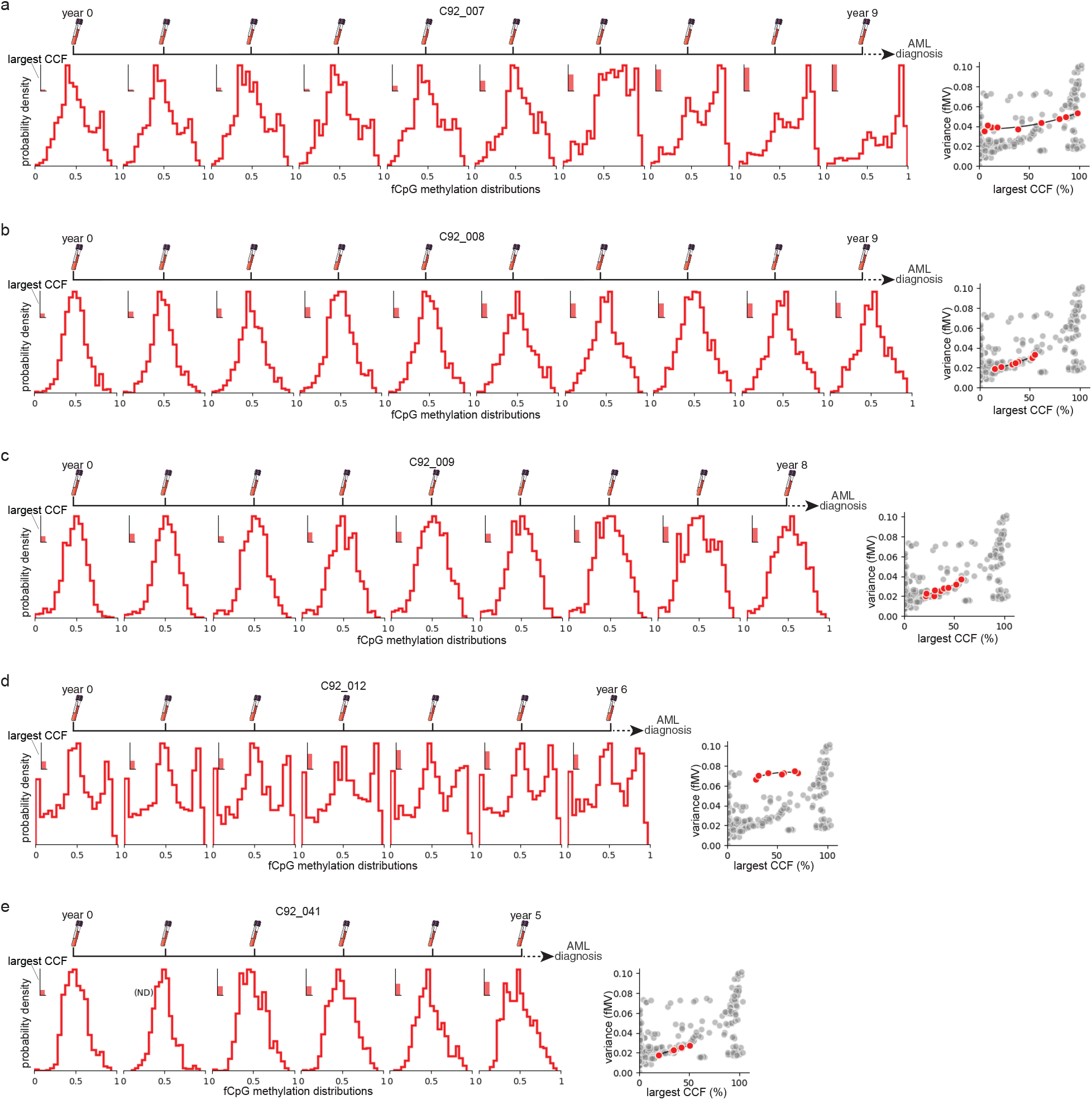
The shape of fCpG methylation distribution is associated with somatic mutation fraction in pre-AML cases. The “W-ness” of the distribution of fCpG methylation fractions, as measured by distribution variance (fMV), in each blood sample increases with clonal burden (i.e., clonal cell fraction, CCF, as measured by the variant allele fraction of somatic mutations). **a-e**. fCpG methylation distributions and clonal burden in the years preceding AML diagnosis for 5 cases with clonal expansions measured by mutant allele fraction (C92_008, C92_009, C92_012, and C92_41). Left panels show the distribution of methylation fractions for 900-1,045 fCpGs in each blood sample collected. Inset bar plots show the clonal burden (from somatic mutational fraction) as the percent of the largest CCF. Right panels show the relationship between fMV and largest CCF for each time point (red dots), a polynomial regression fit (black line), and all other libraries (grey dots). Note that cases C92_007 and C92_012 (**a** and **d**) have higher than expected initial fMVs (reflecting a wider distribution of fCpG methylation fractions) which suggests pre-existing clonal expansions that were not detected by somatic mutations.

**Extended Data Figure 14.**
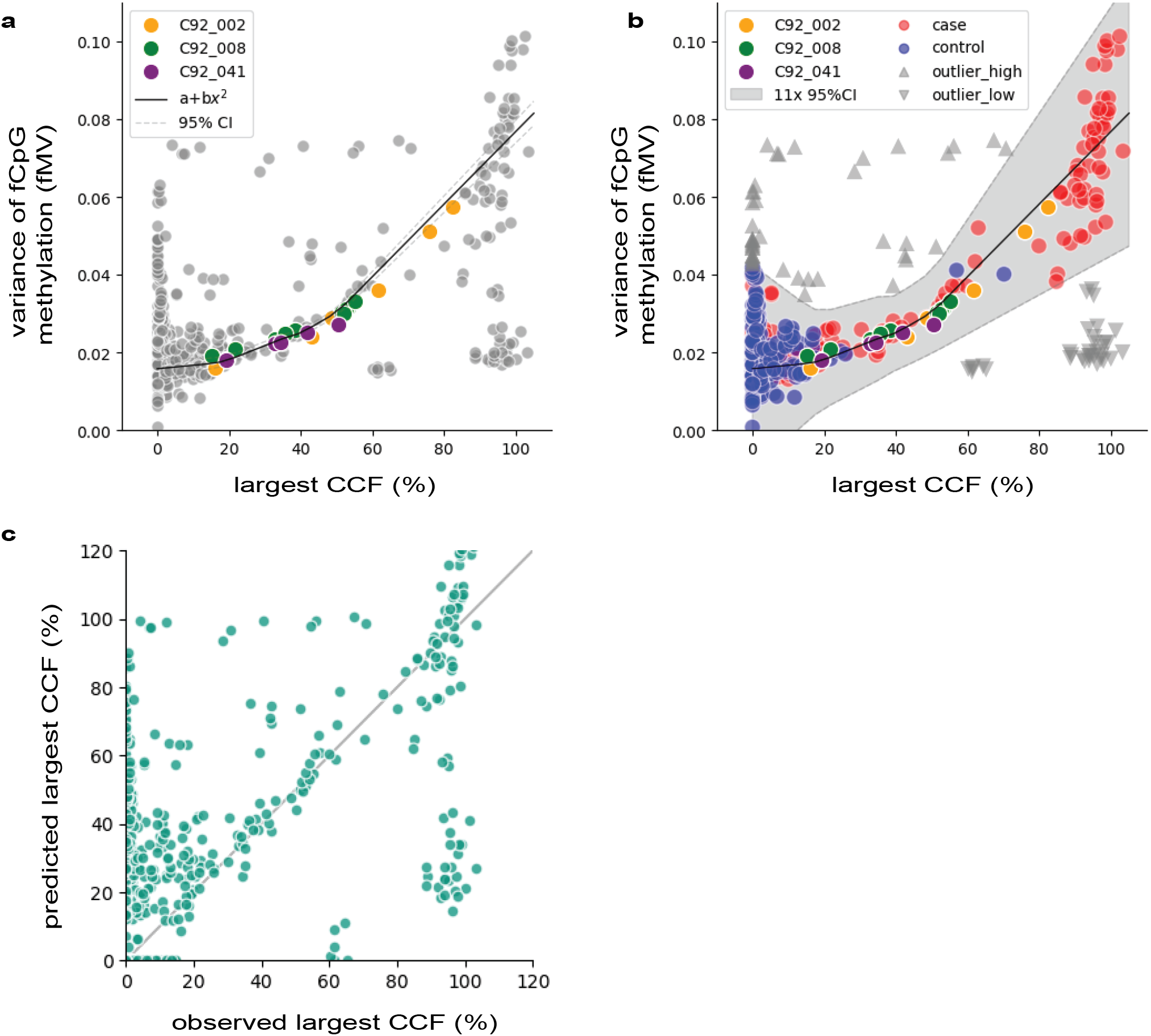
Clonal expansions with undetected drivers and germline mutations miscalled as somatic are revealed using second-order polynomial regression of clonal cell fraction and fCpG methylation in three pre-AML cases. The relationship between the variance of the distribution of fCpG methylation fractions (fMV) and the largest clonal cell fraction (CCF, as measured by somatic mutational abundance) for each blood draw was modelled (**a**) using a second-order polynomial regression of the combined data from pre-AML cases, C92_002, C92_008, and C92_041 (orange, green, and purple, respectively, all other libraries are in grey, R2=0.974). **b** 11-fold expansion of the 95% confidence intervals (grey shading) covered *∼*85% of the data (cases in red, control in blue). Libraries falling outside of this region (grey triangles) are predicted to have clonal expansions with undetected driver mutations (△) or germline mutations miscalled as somatic (▽). **c** Relationship between the predicted clonal burden using the model in (b) and the observed clonal burden (from mutational abundance). Large deviations from y=x are predicted to result from (i) insufficient detection of somatic clonal markers (i.e., missing driver mutations), (ii) germline mutations incorrectly called as somatic (i.e., miscalled germline), or (iii) very early somatic sweeps (>20-30 years prior to sampling) in which synchronized (i.e., clonal) fCpGs have had ample time to epi-mutate and become desynchronized.

**Extended Data Figure 15.**
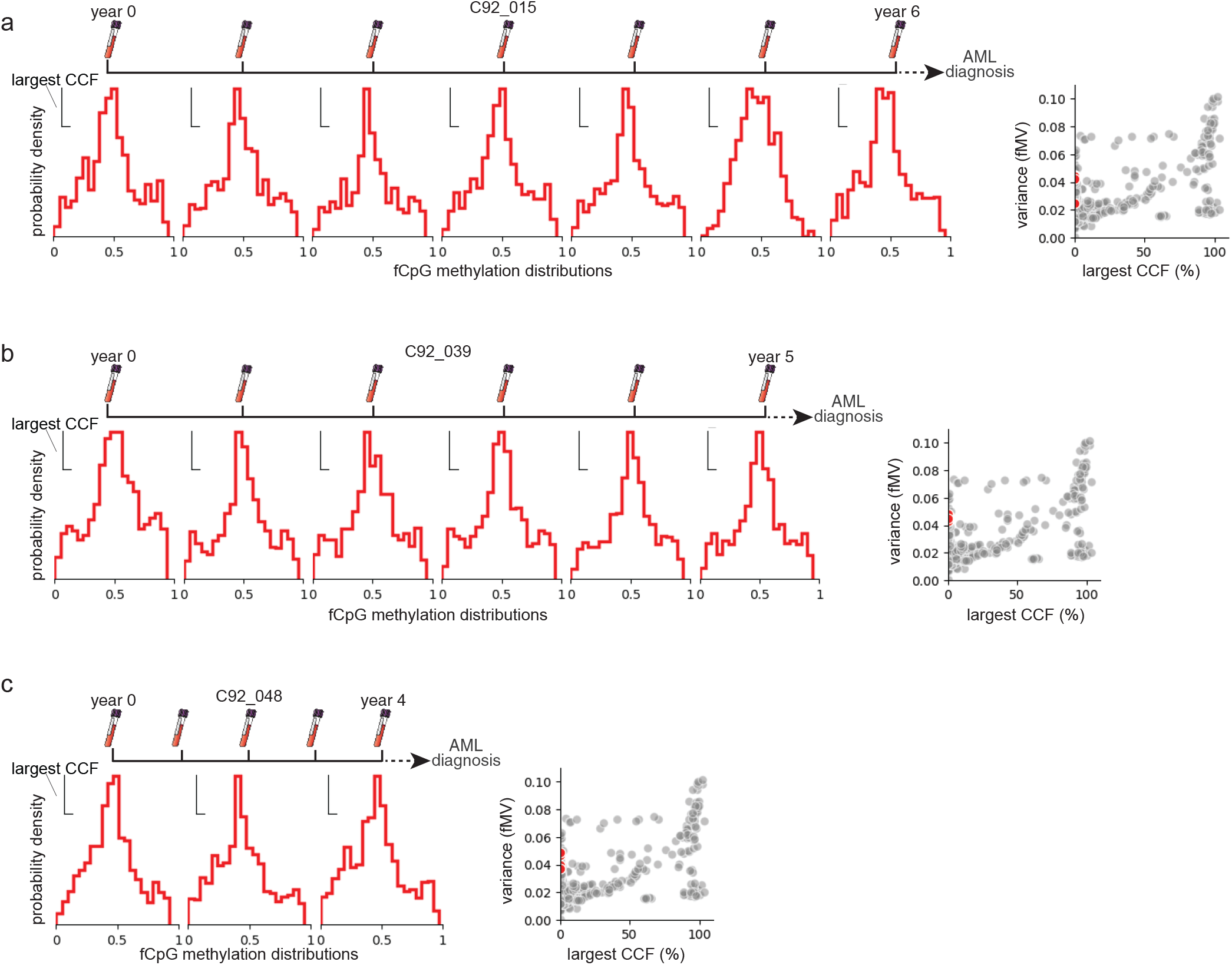
W-shaped fCpG methylation distribution suggests undetected clonal expansions and driver mutations. **a-c**. Distributions of fCpG methylation fractions of three cases (C92_015, C92_039, and C92_048) have high variance (fMV) and are W-shaped, suggesting pre-AML clonal expansions were present in these cases that had no detectable somatic mutations in our comprehensive genetic sequencing panel. Left panels show the distribution of methylation fractions for 900-1045 fCpGs in each blood sample collected. Inset bar plots show the clonal burden (from somatic mutational fraction) as the percent of the largest CCF (*∼*0% for all cases and time points shown here). Right panels show the relationship between fMV and largest CCF for each time point (red dots), a polynomial regression fit (black line), and all other libraries (grey dots). Note that C92_048 libraries had insufficient fCpG coverage in 2 of 5 blood draws (“year 1” and “year 3”) and were omitted.

